# Chromatin remodeling protein CHD4 regulates axon guidance of spiral ganglion neurons in the developing cochlea

**DOI:** 10.1101/2024.01.31.578202

**Authors:** Jihyun Kim, Edward Martinez, Jingyun Qiu, Alejandra Laureano, Anna Maria Hinman, Julie Zhouli Ni, Kelvin Y. Kwan

## Abstract

Spiral ganglion neurons (SGNs) are the primary afferent neurons that convey auditory information from the cochlea to the central nervous system. Pathogenic variants of human chromodomain helicase DNA-binding protein 4 (CHD4) cause Sifrim-Hitz-Weiss (SIHIWES) syndrome, a neurodevelopmental disorder associated with hearing loss. Here, we identified expression of CHD4, an ATP-dependent chromatin remodeler, in developing SGNs and proceeded to delete *Chd4* in mice. Auditory brainstem recordings showed that animals lacking Chd4 in SGNs exhibit hearing loss. SGNs are classified as type I and type II neurons, which display distinct peripheral innervation patterns. Notably, loss of CHD4 resulted in abnormal fasciculation of type I neurons and improper pathfinding of type II fibers. To uncover the underlying molecular mechanisms, we mapped genome-wide CHD4 chromatin occupancy in immortalized multipotent otic progenitor (iMOP)-derived neurons. Gene ontology analysis of CHD4 target genes revealed pathways governing axon guidance, axonal fasciculation, and ephrin receptor signaling. *In vivo*, we observed upregulation of specific Eph/ephrin genes in SGNs from *Chd4*-conditional knockout cochleae. Together, these results demonstrate that CHD4 modulates chromatin at cis-regulatory elements to repress a subset of axon guidance molecules during development. The coordinated regulation of multiple guidance cues is essential for establishing precise neural circuitry in the cochlea. Our findings reveal a critical role for CHD4-dependent epigenetic regulation in neural circuit wiring and offer a translational framework for strategies to regenerate SGNs.

## Introduction

Spiral ganglion neurons (SGNs) in the cochlea are essential for conveying sound information from the cochlear hair cells to the brainstem (Dabdoub, Fritzsch et al. 2016). Assembly of these specialized neural circuits requires distinct developmental steps and activity-dependent synaptic refinement (Sitko and Goodrich 2021). SGNs are categorized as type I and type II neurons. Type I neurons are large, myelinated, and comprise ∼95% of the SGNs that form a single synapse with inner hair cells of the cochlea. Type II neurons are small, unmyelinated neurons that constitute ∼5% of SGNs. Each type II SGN contacts multiple outer hair cells to form *en passant* synapses (Liberman, Wang et al. 2011, Weisz, Lehar et al. 2012, Moser, Karagulyan et al. 2023). The innervation pattern of type I and type II SGNs occurs through a series of defined stages. During early development, neurosensory precursors expressing *Neurog1*, a key transcription factor required for SGN development, give rise to SGNs. Starting around embryonic day 10 (E10), the *Neurog1*-expressing precursors delaminate from the otic vesicle by E11, forming neuronal precursors. Between E12-14, SGNs extend their peripheral axons towards the cochlear hair cells and central axons towards the brainstem. By E15, initial innervation patterns are established between SGNs, their target hair cells, and the cochlear nucleus in the brainstem. Starting at E16, the neural circuits undergo refinement through activity-dependent mechanisms, including synapse formation and refinement (Appler and Goodrich 2011, Coate and Kelley 2013). The initial establishment of circuit assembly in SGNs requires cellular processes such as axon guidance and repulsion cues to pattern and assemble the circuits (Sitko and Goodrich 2021). However, how epigenetic mechanisms regulate the timing and expression of axon guidance genes during SGN development remains poorly understood.

Epigenetic changes can significantly influence the expression of axon guidance genes during development. Changes such as DNA methylation and histone modifications control gene expression patterns and the timing of guidance cues. For instance, epigenetic modifications can regulate the expression of genes encoding guidance molecules like netrins, ephrins, slits, semaphorins, and neuropilin, which are essential for directing peripheral axon patterns (Gillespie, Marzella et al. 2005, Coate, Raft et al. 2012, Defourny, Poirrier et al. 2013, Coate, Spita et al. 2015, Kim, Ibrahim et al. 2016, Salehi, Ge et al. 2017). By modulating access to cis-regulatory elements that control the transcription of these genes, epigenetic changes can regulate the timing and expression levels of axon guidance genes, thereby controlling wiring.

Mutations in genes encoding epigenetic modifiers implicated in syndromic hearing loss implicate these molecules in cochlear development. Pathogenic variants in DNA methyltransferases (DNMT1), histone methyltransferases (EHMT1, KMT2D), acetyltransferase (KAT6B), and the chromodomain helicase DNA-binding (CHD) protein family are associated with hearing loss (Layman and Zuo 2014). The CHD family of proteins mobilizes and rearranges nucleosomes to alter chromatin accessibility. Pathogenic variants in CHD4 and CHD7 are associated with Sifrim-Hitz-Weiss and CHARGE syndrome, respectively (Micucci, Sperry et al. 2015, Clapier, Iwasa et al. 2017). Sifrim-Hitz-Weiss (SIHIWES) is an autosomal dominant intellectual developmental disorder with variable congenital abnormalities. Patients with SIHIWES show delayed development, intellectual disability, facial dysmorphism, and ear abnormalities, including hearing loss (Sifrim, Hitz et al. 2016, Weiss, Terhal et al. 2016).

However, the contribution of CHD4 to specific inner ear cell types remains unclear. In the mouse, the expression of CHD4 and components of the nucleosome remodeling and deacetylase (NuRD) complex are detected in many cell types of the cochlear sensory epithelium (Layman, Sauceda et al. 2013). CHD4 has been studied in the context of the NuRD complex (Tong, Hassig et al. 1998, Xue, Wong et al. 1998, Zhang, LeRoy et al. 1998). The NuRD complex contains CHD4, histone deacetylase (HDAC1 and HDAC2), and additional subunits that regulate chromatin accessibility and transcription. The NuRD complex localizes to cis-regulatory elements to affect gene expression (Morra, Lee et al. 2012, Watson, Mahajan et al. 2012, Marques, Gryder et al. 2020).

A decrease in transcription caused by the NuRD complex affects a myriad of cellular processes, including maintenance of cell identity, DNA damage repair, cell cycle progression, and cancer (Polo, Kaidi et al. 2010, Hung, Kohnken et al. 2012, Pan, Hsieh et al. 2012, Hosokawa, Tanaka et al. 2013, Yang, Yamada et al. 2016, Xia, Huang et al. 2017, Bornelov, Reynolds et al. 2018). CHD4 binds to thousands of sites in the mammalian genome in a cell-type-specific manner, and studies show that CHD4 forms distinct protein complexes with NuRD-independent functions (Low, Webb et al. 2016, Hoffmeister, Fuchs et al. 2017, Kaaij, Mohn et al. 2019).

To define the function of CHD4 in SGN development, we employed a conditional knockout (cKO) model to delete *Chd4* in neurosensory progenitors and examine its effects on SGN wiring. We examined CHD4 function in SGNs using a transgenic mouse line harboring the *Neurog1* (*Ngn1*) CreER^T2^ (Koundakjian, Appler et al. 2007), *Chd4* cKO alleles (Williams, Naito et al. 2004), and a tdTomato reporter to label SGN lineages (Madisen, Zwingman et al. 2010).

## Materials and Methods

### Generation of *Chd4* cKO mice

All procedures were conducted in accordance with institutional animal care guidelines and committee-approved research protocols at Rutgers University. *Chd4^tm1Kge/tm1Kge^* (*Chd4* floxed) animals (RRID: MGI:3641408) were obtained (Williams, Naito et al. 2004). *Neurog1* (*Ngn1*) CreER^T2^ animals and Ai9 tdTomato reporter animals were purchased from Jackson Laboratory (RRID: IMSR_JAX:008529 and RRID: IMSR_JAX:007909, respectively) (Koundakjian, Appler et al. 2007, Madisen, Zwingman et al. 2010). PCR genotyping was performed using the EconoTaq Plus Green 2X Master Mix (Biosearch Technologies, #30033-1) and primer pairs for wild-type and *Chd4* floxed alleles (Table 1). For Cre-induced excision of loxP-flanked exons that code for the ATPase domain of *Chd4*, a mixture of tamoxifen and β-estradiol was diluted in corn oil. β-estradiol was included to help alleviate the anti-estrogen effects of tamoxifen treatments. Tamoxifen doses for pregnant female mice were adjusted for maternal body weight. Pregnant dams were gavaged with 6.25mg/kg of tamoxifen (Sigma Aldrich, #T5648) and 6.25µg/kg of β-estradiol (Sigma Aldrich, #E8875) daily on embryonic days (E) 8.5- 10.5 to induce Cre activity. For staging timed embryos, the morning that vaginal plugs were observed, pregnant female mice were considered to be carrying E0.5 embryos.

**Table 1.**
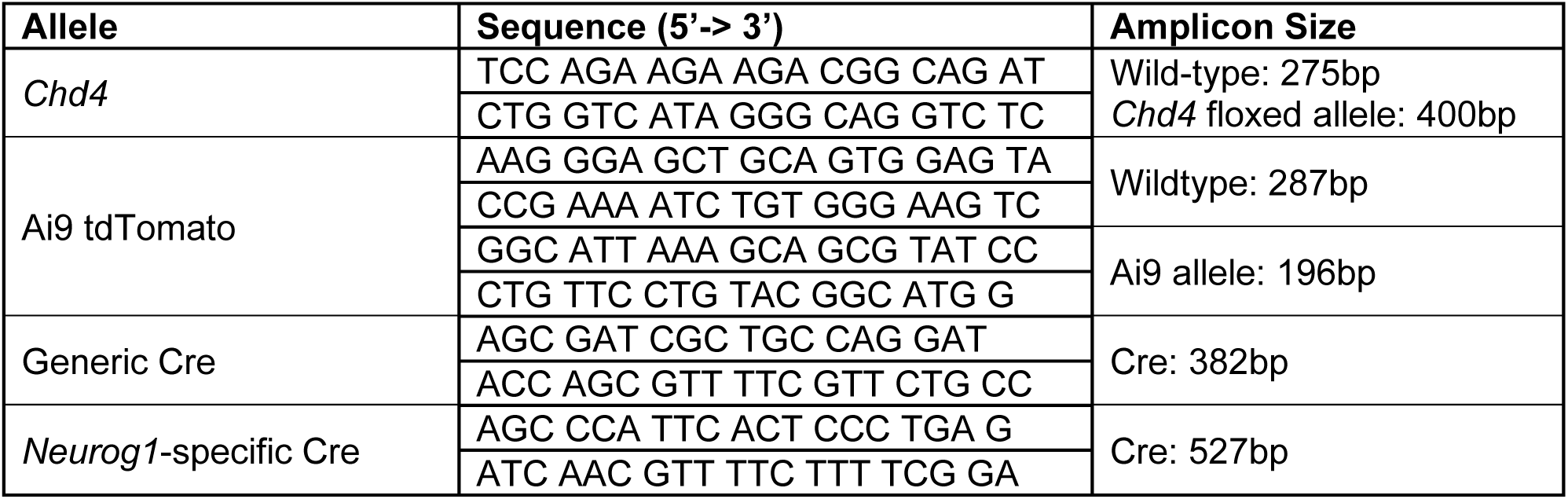
List of primers used for genotyping.

### Auditory Brainstem Recordings

Auditory brainstem recordings (ABRs) were obtained from 3-4-week-old mice anesthetized with 75 mg/kg of ketamine and 15 mg/kg of xylazine (Henry Schein). Platinum needle electrodes (Grass Telefactor) were placed subdermally at the vertex (active) and mastoid (reference), with a ground electrode positioned near the tail. Stimulus generation and signal acquisition were performed using Tucker-Davis System 3 hardware controlled by BioSigRZ software (Tucker-Davis Technologies). ABRs were evoked using click stimuli and tone pips (4 to 32 kHz; 5 ms duration; 0.5 ms rise/fall time) delivered through a closed field speaker. Stimuli were presented at a rate of 25 s^-1^. Sound pressure levels (SPL) were decreased in 10 dB steps from 90 to 10 dB SPL to determine thresholds. Responses were amplified, filtered, and averaged over repeated presentations. Data from recordings were exported to CSV files and analyzed using the Auditory Brainstem Response (ABR) Waveform Analysis program (Burke et al., 2023). Thresholds were defined as the lowest SPL that reproducibly generates identifiable waveforms. Wave 1 amplitudes and latencies were measured from suprathreshold responses. Wave 1 amplitude was defined as the peak-to-trough differences, and latency was defined as the time from stimulus onset to the wave 1 peak (Sergeyenko, Lall et al. 2013).

### Tissue fixation and preparation

Embryos or postnatal pups were sacrificed for inner ear dissection. The temporal bone was identified and removed after bisecting the mouse head and removing the brain. The inner ear was separated from the temporal bone, fixed directly in 4% formaldehyde in 1X PBS overnight at 4°C, and washed with 1X PBS overnight at 4°C. Extraneous tissue and bone surrounding the cochlea were removed to obtain the cochlear epithelium and the cochlear duct. Reissner’s membrane was removed to expose the sensory epithelium, which was then processed for immunostaining and *in situ* hybridization. After processing, the cochleae, including the spiral ganglion and sensory epithelium, were flat-mounted.

### Immunohistochemistry

The micro-dissected cochlea tissues fixed in 4% formaldehyde in 1X PBS were permeabilized in wash buffer (1X PBS with 0.1% Triton X-100) for 10 min before being incubated for at least 1 hour in blocking buffer (1X PBS with 0.1% Triton X-100 and 10% normal goat serum) at room temperature. Samples were incubated overnight at 4°C with appropriate primary antibodies diluted in the blocking solution. Samples were then washed with 1X PBS and incubated with the appropriate combinations of DAPI (1 µg/ml) and Alexa Fluor 488-, Alexa Fluor 568-, or Alexa Fluor 647-conjugated secondary antibodies (Thermo Fisher Scientific) for 2 hours at room temperature and washed in 1X PBS before mounting in Prolonged Gold Antifade (Invitrogen, #P36934).

iMOP cells were fixed in 4% formaldehyde in 1X PBS for 20 minutes at room temperature, incubated in blocking buffer for 1 hour at RT, and incubated overnight with the primary antibodies in blocking buffer at 4°C. The next day, cells were washed with 1X PBS and incubated with appropriate combinations of DAPI (1 µg/ml) and Alexa Fluor 488-, Alexa Fluor 568-, or Alexa Fluor 647-conjugated secondary antibodies (Thermo Fisher Scientific) in a blocking buffer for 2 hours at RT. Samples were washed with 1X PBS and mounted in Prolonged Gold Antifade. All antibodies used for immunostaining are listed in Table 2.

**Table 2.**
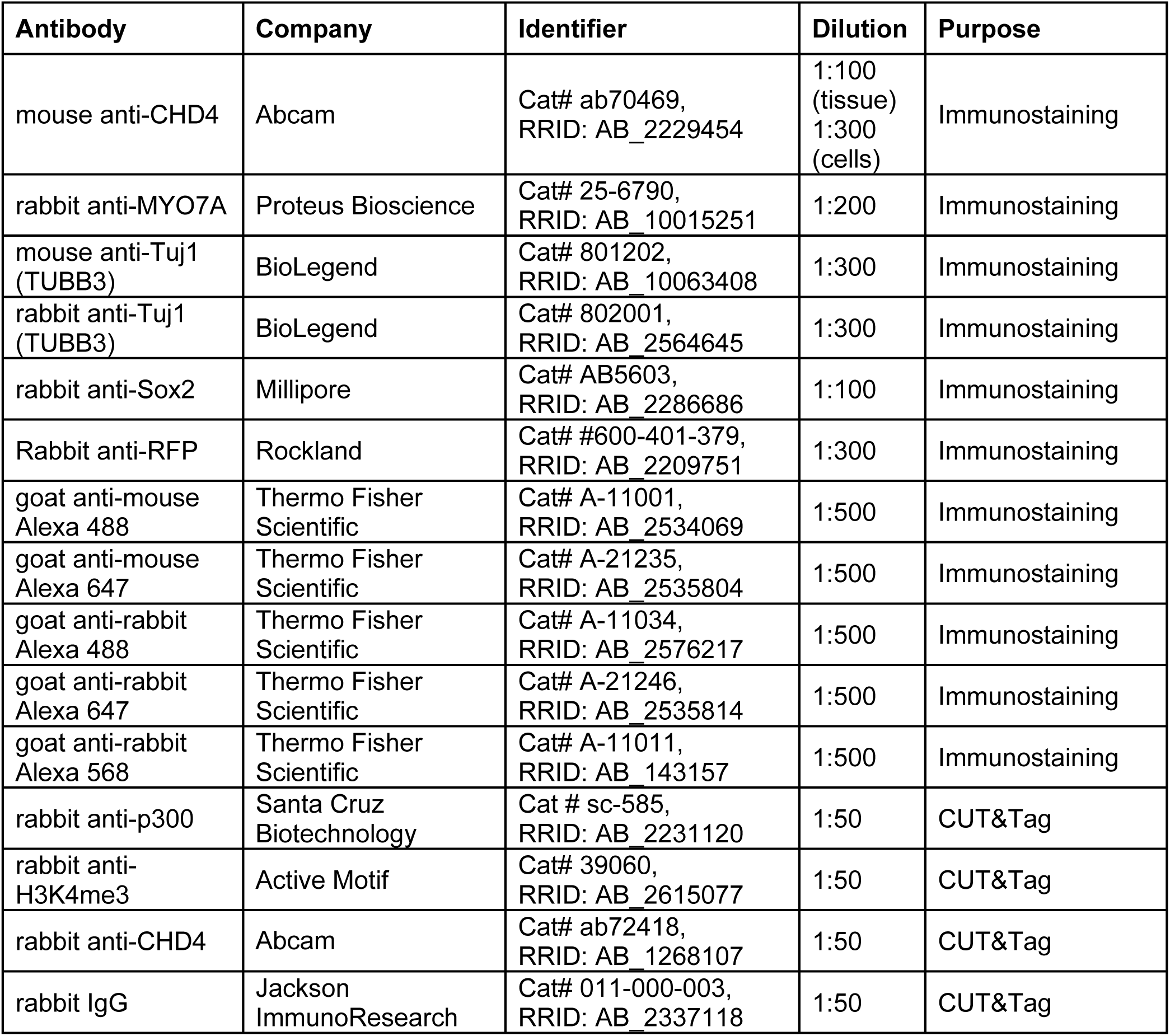
List of antibodies used for immunostaining and CUT&Tag.

| **Antibody** | **Company** | **Identifier** | **Dilution** | **Purpose** |
| --- | --- | --- | --- | --- |
| mouse anti-CHD4 | Abcam | Cat# ab70469,<br>RRID: AB_2229454 | 1:100<br>(tissue)<br>1:300<br>(cells) | Immunostaining |
| rabbit anti-MYO7A | Proteus Bioscience | Cat# 25-6790,<br>RRID: AB_10015251 | 1:200 | Immunostaining |
| mouse anti-Tuj1<br>(TUBB3) | BioLegend | Cat# 801202,<br>RRID: AB_10063408 | 1:300 | Immunostaining |
| rabbit anti-Tuj1<br>(TUBB3) | BioLegend | Cat# 802001,<br>RRID: AB_2564645 | 1:300 | Immunostaining |
| rabbit anti-Sox2 | Millipore | Cat# AB5603,<br>RRID: AB_2286686 | 1:100 | Immunostaining |
| Rabbit anti-RFP | Rockland | Cat# #600-401-379,<br>RRID: AB_2209751 | 1:300 | Immunostaining |
| goat anti-mouse<br>Alexa 488 | Thermo Fisher<br>Scientific | Cat# A-11001,<br>RRID: AB_2534069 | 1:500 | Immunostaining |
| goat anti-mouse<br>Alexa 647 | Thermo Fisher<br>Scientific | Cat# A-21235,<br>RRID: AB_2535804 | 1:500 | Immunostaining |
| goat anti-rabbit<br>Alexa 488 | Thermo Fisher<br>Scientific | Cat# A-11034,<br>RRID: AB_2576217 | 1:500 | Immunostaining |
| goat anti-rabbit<br>Alexa 647 | Thermo Fisher<br>Scientific | Cat# A-21246,<br>RRID: AB_2535814 | 1:500 | Immunostaining |
| goat anti-rabbit<br>Alexa 568 | Thermo Fisher<br>Scientific | Cat# A-11011,<br>RRID: AB_143157 | 1:500 | Immunostaining |
| rabbit anti-p300 | Santa Cruz<br>Biotechnology | Cat # sc-585,<br>RRID: AB_2231120 | 1:50 | CUT&Tag |
| rabbit anti-H3K4me3 | Active Motif | Cat# 39060,<br>RRID: AB_2615077 | 1:50 | CUT&Tag |
| rabbit anti-CHD4 | Abcam | Cat# ab72418,<br>RRID: AB_1268107 | 1:50 | CUT&Tag |
| rabbit IgG | Jackson<br>ImmunoResearch | Cat# 011-000-003,<br>RRID: AB_2337118 | 1:50 | CUT&Tag |

### Single-molecule *in situ* hybridization

Single-molecule fluorescence in situ hybridization (smFISH) was performed using RNAscope (Wang et al., 2012). The RNAscope Multiplex Fluorescent Reagent Kit v2 (Advanced Cell Diagnostics, #323100) was used to detect target RNAs according to the manufacturer’s instructions. RNAscope probes are listed in Table 3. Immunohistochemistry was performed after the smFISH protocol to enhance and detect tdTomato protein. Samples were incubated in blocking buffer (1X PBS, 10% goat serum, and 0.3% Triton X-100) for 1 hour at room temperature, followed by overnight incubation with rabbit anti-RFP in blocking buffer at 4°C. After incubation with primary antibodies, samples were washed with 1x PBS and then incubated with appropriate combinations of DAPI (1 µg/ml) and Alexa Fluor 568-conjugated secondary antibodies (Thermo Fisher Scientific) in a blocking buffer for 2 hours at room temperature. Samples were washed with 1X PBS and mounted in Prolonged Gold Antifade. All antibodies used with RNAscope are listed in Table 2.

**Table 3.** List of RNAscope probes used for smFISH.

| **RNAscope Probe** | **Accession Number** | **Probe Region** | **Catalog Number** | **Company** |
| --- | --- | --- | --- | --- |
| Mm-Calb2-C2 | NM_007586.1 | 147-1265 | 313641-C2 | Advanced Cell<br>Diagnostics, Inc. |
| Mm-Epha4 | NM_007936.3 | 291-1191 | 419081 |  |
| Mm-Calb2-C3 | NM_007586.1 | 147-1265 | 313641-C3 |  |
| Mm-Epha7-C2 | NM_010141.4 | 1024-1981 | 430961-C2 |  |
| Mm-Efnb2 | NM_010111.5 | 174-1316 | 477671 |  |

### Fluorescence micrograph acquisition and visualization

Samples were mounted on a #1.5 cover glass and imaged using a Zeiss LSM800 point-scanning confocal microscope. Antibody-conjugated fluorophores and tdTomato fluorescent protein were excited using four laser lines (405, 488, 561, and 633nm). 1024X1024 images were acquired using either the 40X (1.4 NA) or 63X (1.4 NA) objective with 1.5-2x digital zoom and 4X Kalmann averaging. Image brightness and contrast were adjusted and viewed using Zen Blue (Zeiss) or ImageJ.

### Cell Culture for iMOP cells

Proliferating iMOP cells were cultured in suspension with DMEM/F12 (Gibco, #11320033) containing N21-MAX supplement (R&D Systems, #AR008), 25 µg/ml carbenicillin (Fisher Scientific, # BP2648-1), and 20 ng/ml bFGF (PeproTech, #450-33). Cells were passaged weekly by dissociation using TrypLE Express (Gibco, #12604013), harvested by centrifugation, and resuspended in fresh medium. To generate iMOP-derived neurons, tissue culture dishes or 12mm coverslips placed in a 24-well plate were coated with 10 µg/mL of poly-D-lysine (PDL; Corning, #354210) for 1 hour at 37°C, then coated overnight with 10 µg/mL of laminin (Gibco, # 23017015) at 37°C, and washed 3 times with 1X PBS before plating cells. Proliferating iMOP cultures were dissociated in TrypLE Express, harvested by centrifugation, and resuspended in neurobasal media (Gibco, #2113049). Resuspended cells were counted using a Moxi counter, and particles between 9-13 µm were considered as cells. Dissociated iMOP cells were plated in PDL-laminin-coated culture dishes (1.5-2 x 10^5^ cells /dish for CUT&Tag) or coverslips in a 24-well plate (0.8 x 10^5^ cells/well for immunostaining) and cultured in a neurobasal medium containing N21-MAX, 2 mM L-glutamine (Gibco, #25030081), and 1 µM K03861 (Selleck chemicals, #S8100) for neuronal differentiation. On day 3 of culture, the Culture One Supplement (Gibco, A3320201) was added to the media. Fresh medium was added every other day, and neuronal cultures were harvested 7 days after being plated for neuronal differentiation and then used for CUT&Tag or immunostaining.

### Cleavage Under Targets and Tagmentation

Cells from proliferating iMOPs and iMOP-derived neurons were harvested and processed using the CUT&Tag-IT Assay Kit (Active Motif, 53160) according to the manufacturer’s instructions. Cells were gently dissociated using TrypLE Express, washed with 1x PBS, harvested by centrifugation, and counted. For each CUT&Tag reaction, 500,000 cells were used. The supernatant from the cell pellet was removed, and cells were resuspended in a wash buffer and placed on ice. Cells were enriched using magnetic Concanavalin A beads and resuspended in an antibody buffer containing a primary antibody to the protein of interest. H3K4me3, p300, CHD4, and IgG antibodies were used for individual samples. All antibodies are listed in Table 2. Two independent samples were used for each antibody condition. Primary antibodies were incubated with the cells for 2 hours at room temperature or overnight at 4°C. After incubation, the cells were washed and incubated in a buffer containing the assembled pA-Tn5 transposomes and incubated for 1 hour at RT. Cells were washed and processed for tagmentation by incubation in the appropriate buffer at 37 °C for 1 hour. A solution containing EDTA, SDS, and proteinase K was added to stop the reaction and solubilize DNA fragments. The solution was placed on a magnetic stand, and the supernatant containing soluble DNA fragments was retained. The DNA solution was purified using a column. The DNA fragments were subjected to PCR amplification and purified using SPRI beads. Libraries generated were quantified and subjected to quality control before being sequenced on an Illumina HiSeq 3000 platform (2x150 bp). All antibodies used for CUT&Tag are listed in Table 2.

### Peak calling

Paired-end CUT&Tag reads were aligned to the mm10 genome assembly with bowtie2. Only mapped reads were retained, written to the output, and used for the downstream analysis (Langmead and Salzberg 2012). The bowtie2 SAM output files were converted into BAM files using Samtools (Li, Handsaker et al. 2009). After alignment, sequence reads were scaled using the bamCoverage command in deepTools to remove duplicates and generate normalized bigwig files with reads as Reads Per Kilobase per Million (RPKM) (Ramirez, Dundar et al. 2014). The UCSC bigWigToBedGraph utility was used to generate bedgraph files from bigwig files (Kent, Zweig et al. 2010). Peaks were called using Sparse Enrichment Analysis for CUT&RUN (SEACR) (Meers, Tenenbaum et al. 2019). SEACR output bed files from CHD4, H3K4me3, and p300 were generated using CHD4, H3K4me3, and p300 bedgraph files as target files, while using the IgG bedgraph file as a threshold. BEDTools (Quinlan and Hall 2010) was used to identify consensus peaks with at least 10% reciprocal overlap between replicate samples. Only consensus peaks were used for downstream analysis. CUT&Tag signal tracks were visualized on Integrative Genomic Viewer (IGV) (Robinson, Thorvaldsdottir et al. 2011).

### Identifying CHD4-bound promoters and enhancers

To perform a global survey of CHD4-bound promoters and enhancers, we first identified them in proliferating iMOPs and iMOP-derived neurons. H3K4me3 and p300 were used as promoter and enhancer marks, respectively. CHD4-bound promoters (CHD4+ H3K4me3+) and enhancers (CHD4+ p300+) in iMOPs were identified using the BEDTools intersect command. CHD4+ H3K4me3+ regions were identified as those with at least one base-pair overlap between CHD4 and H3K4me3 peaks. CHD4+ p300+ regions were defined using the same strategy. SEACR total signals were obtained from the SEACR output file and plotted as box-and-whisker plots to show the quartiles, mean (circle), and median (line). Cell state-specific CHD4-bound promoters or enhancers were identified based on their presence or absence in progenitors and neurons. Genome arithmetic was done using bedtools.

### Heatmap and profile plots

Heatmaps and profile plots were generated using deepTools. The signal and region files were obtained from the CUT&Tag data. Normalized bigwig files from two replicates were merged using bigwigCompare to acquire average bigwig files and used as signal files. The summit regions corresponding to the maximum SEACR peak signal from CHD4 were used as region files. DeepTools was used to visualize CHD4 and H3K4me3 reads as heatmaps or profile plots. Plots were centered at the summits of CHD4+ H3K4me3+, and the average CHD4 or H3K4me3 bigwig signals were plotted ± 3 kb from the summits. To visualize CHD4 and p300 reads, the summits from CHD4+ p300+ were used as the center, and the average bigwig signals of either CHD4 or p300 bigwig files were plotted within ± 3kb of the summits.

### Quantification of heatmap total signals

To quantify CUT&Tag signals displayed in heatmaps, the -outFileNameMatrix option in deepTools was used to retrieve the matrix of signal values underlying the heatmaps. CUT&Tag signals within specified genomic regions were added to obtain the summed signal values. Heatmap signals between progenitors and neurons within the ±3 kb window were visualized as box-and-whisker plots.

### Gene ontology (GO) analysis

To annotate CHD4+ common and neuron-specific promoters and enhancers, ChIPpeakAnno was used to identify the closest genes and their corresponding Ensembl gene IDs for those regions (Zhu, Gazin et al. 2010). Unique Ensembl gene IDs near CHD4-bound promoters and enhancers were used for GO analysis with DAVID (Database for Annotation, Visualization and Integrated Discovery) (Huang, Sherman et al. 2009).

### *De Novo* Motif Discovery, Comparison, and Visualization

Genomic coordinates from SEACR-called CHD4 summits were used to define regions of interest. The regions were converted to GRanges objects. Corresponding DNA sequences were extracted from the *Mus musculus* (mm10) reference genome using the BSgenome.Mmusculus.UCSC.mm10 package. The MEME suite was employed for motif analysis using the memes, an R package (Bailey, Johnson et al. 2015, Nystrom and McKay 2021). MEME (Multiple Expectation maximizations for Motif Elicitation) was used for *de novo* motif discovery. MEME was configured to identify 20 motifs using the anr (Any Number of Repetitions) model, with motif widths ranging from 6-15 base pairs, searching both forward and reverse complement strands. Position frequency matrices (PFMs) of discovered motifs were normalized to probabilities and visualized as sequence logos using the universal motif and ggplot2 packages.

Discovered *de novo* motifs were compared against known transcription factor binding motifs from the JASPAR2024 CORE database for *Mus musculus* using TomTom. *De novo* motifs, corresponding to the most statistically significant match in JASPAR2024, were based on e-values < 1 × 10^-5^. The best-matching transcription factor names were used for motif annotation. To assess motif occurrence and distribution around the transcriptional start site (TSS), FIMO (Find Individual Motif Occurrences) was utilized. The regions ±1kb around the TSS of axon guidance genes harboring CHD4 summits were extracted. FIMO was used to scan these regions for occurrences of all *de novo* motifs. FIMO results were filtered to retain only statistically significant matches with a p-value < 0.05. Motif occurrences with best-match transcription factor names using the TomTom results were mapped and visualized with each axon guidance gene using ggplot2. Individual motif instances were visualized as points relative to the TSS.

### Statistical analysis

For normally distributed data, statistical analyses were performed using either R (R version 4.4.3) or OriginPro (OriginLab). Data are presented as mean ± standard error of the mean (SEM) and were subjected to an unpaired, two-tailed Student’s *t*-test to determine statistical significance. P-values less than 0.05 were considered statistically significant. Unless noted, significance levels are defined as *p < 0.05, **p < 0.01, and ***p < 0.001. Bar graphs display the means ± SEM overlaid with individual data points, whereas violin plots show the data distribution and individual data points. Values displayed in the text indicated the means ± SEM.

Data for CUT&Tag and RNAscope were found to be non-normally distributed using the Shapiro-Wilk normality test. The median was presented and tested with a nonparametric Wilcoxon rank-sum test to determine whether the paired samples differed statistically. Data analysis was performed in R. The normalized read counts (RPKM) from CUT&Tag were displayed as box-and-whisker plots to show quartiles, means, and medians. Horizontal lines within the box denoted the median values, while the circles represented the means. The number displayed in the text indicates the median ± standard deviation (SD) of CUT&Tag signals for the box-and-whisker plots. The RNAscope images were counted to determine the distribution, mean, and median of puncta counts per cell. Unless noted, the p-values from the Wilcoxon rank-sum test were defined as p < 0.05 to be significant. The numbers displayed indicate the median ± SD in the text and the median of puncta counts in the figure. The variance was calculated as the sum of the squared differences between puncta counts and the sample mean, divided by the sample size minus one. Levene’s test for homogeneity of variances was used to determine whether the variances were statistically significant. The p-values from Levene’s test were considered significant if p < 0.05.

## Results

### CHD4 is expressed in the cochlea

Spiral ganglion neurons (SGNs) are bipolar or pseudounipolar cells whose somas reside within the modiolus of the cochlea. From the modiolus, each SGN extends a peripheral axon through Rosenthal’s canal to innervate sensory hair cells and a central axon to the cochlear nucleus within the brainstem. SGNs are broadly classified as type I and type II neurons. Type I SGNs are the primary afferents that consist of 90-95% of the neuronal population in the spiral ganglion. Type I SGNs are myelinated and form a single synaptic bouton with inner hair cells that reside in the organ of Corti (Fig. 1A). Multiple type I SGNs synapse along the basolateral surface of inner hair cells (IHCs) (Liberman 1982). Type I SGNs differ in sensitivity to sound and spontaneous firing rate (SR), as revealed by single-fiber recordings in the cat auditory nerve (Kiang, Pfeiffer et al. 1965). The relationship between threshold and SR predicted three distinct populations of low-, medium-, and high-SR neurons (Liberman 1978). These three subtypes are present irrespective of tonotopic position along the length of the cochlea (Borg, Engstrom et al. 1988, Schmiedt 1989, Winter, Robertson et al. 1990, el Barbary 1991, Taberner and Liberman 2005, Shrestha, Chia et al. 2018, Sun, Wang et al. 2022). SGNs display different receptors and ion channel regulators that shape their sensitivity and SR. The molecular heterogeneity correlated with the differences in electrophysiological properties of these neurons (Adamson, Reid et al. 2002, Chen, Xue et al. 2011, Liu, Manis et al. 2014, Liu and Davis 2014).

**Figure 1.**
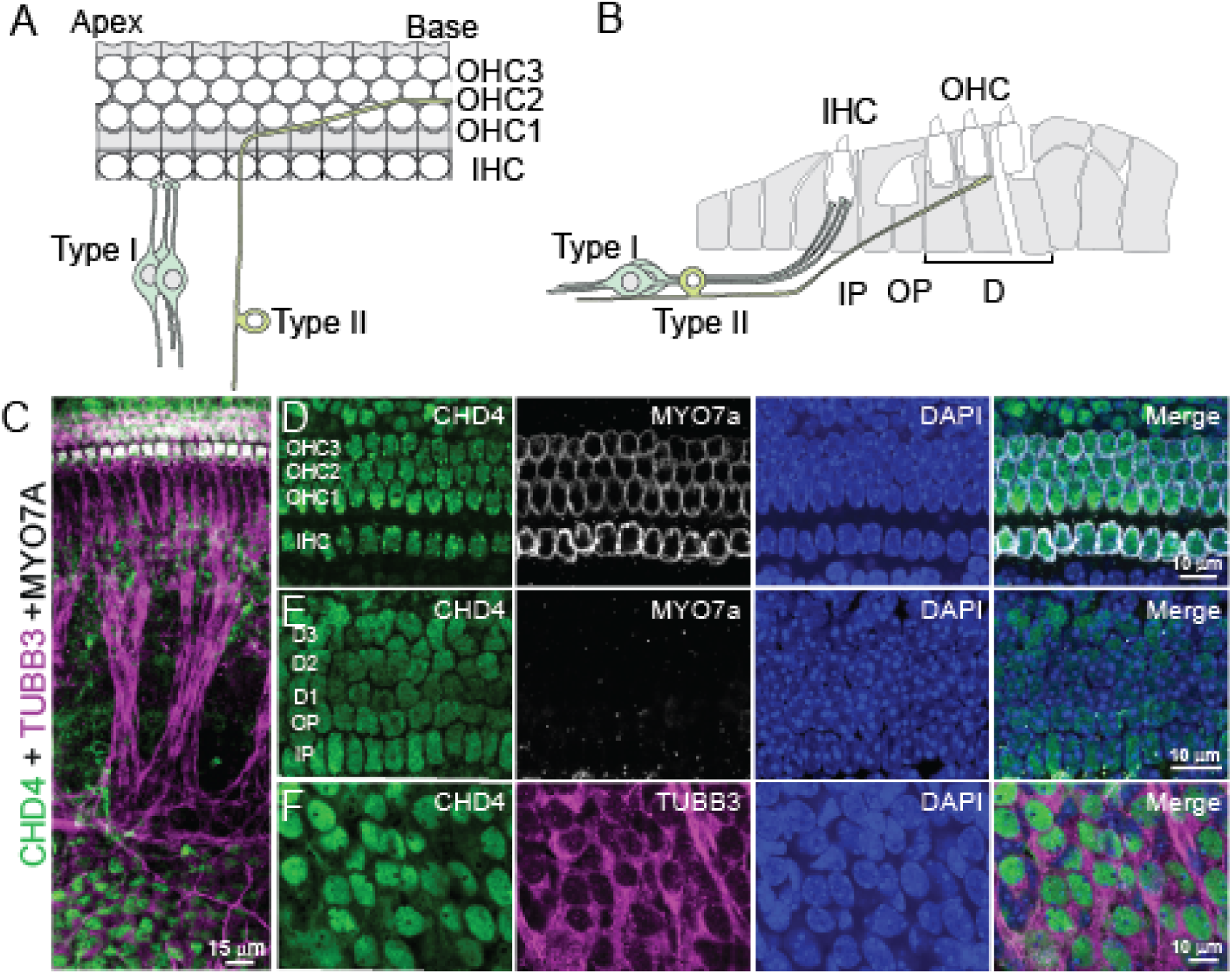
CHD4 expression in the spiral ganglion and the organ of Corti. (A) Top-down schematic depicting the innervation pattern of type I and type II SGNs. Type I SGNs extend single, unbranched neurites that innervate an individual inner hair cell (IHC). Type II SGNs extend neurites into the outer hair cell (OHC) region, turn basally, and can contact multiple OHCs. (B) Sagittal schematic of the innervation pattern. Multiple type I SGNs innervate a single IHC. Type II neurites make a basal turn in the region near Deiters’ cells, a supporting cell type. (C) Immunofluorescence labeling of CHD4 (green), TUBB3 (magenta), and MYO7A (white) from a flat-mounted cochlea. TUBB3 and MYO7A label neurons and hair cells, while DAPI (blue) marks nuclei. Different regions of immunolabeled E18.5 cochlea. (D) Expression of CHD4 in the single row of inner hair cells (IHC) and the three rows of outer hair cells (OHC1-3). (E) CHD4 labeling in supporting cells, including inner pillar (IP), outer pillar (OP), and three rows of Dieter cells (DC1-3). The nuclei of supporting cells reside underneath hair cells. (F) CHD4 labeling in SGNs. Scale bar as marked.

Type II afferent neurons account for 5–10% of the neuronal population (Spoendlin 1972, Ruggero, Santi et al. 1982). Type II SGNs are pseudounipolar and unmyelinated neurons that turn towards the cochlear base (Zhang and Coate 2017). During outgrowth, Type II SGN peripheral axons turn after the growth cone passes between the basolateral surfaces of the inner pillar cells. Turning occurs among the outer pillar and Deiters’ cells as they are directed towards the cochlear base (Fig. 1A). This process is mediated by planar cell polarity complexes formed between cochlear-supporting cells, thereby enabling non-cell-autonomous regulation of axon pathfinding (Ghimire, Ratzan et al. 2018, Ghimire and Deans 2019). The peripheral axons then gradually ascend apically toward the outer hair cells (OHCs) after turning to join other type II axons in one of three outer spiral bundles that extend along the length of the cochlea (Fig. 1B). Each type II afferent neuron innervates multiple OHCs from the same row, with individual OHC receiving an *en-passant* contact from a type II afferent neuron (Zhang and Coate 2017).

To determine the expression of CHD4, neonatal cochleae from embryonic day (E) 18.5 embryos were harvested. At this point, type I and type II SGNs are morphologically distinct, and the neurites have extended toward IHCs and OHCs, respectively. The cochleae were subjected to immunostaining with CHD4 and TUBB3 antibodies to mark the presence of type I and type II SGNs along with CHD4 (Fig. 1C). The fiber tracks from their innervation pattern can identify the type of SGNs. As previously described, many different cell types within the organ of Corti, including IHCs and OHCs, express CHD4 (Layman, Sauceda et al. 2013). We confirmed the presence of CHD4 in MYO7A labeled hair cells (Fig. 1D). CHD4 was also expressed in the nuclei of inner pillar, outer pillar and the three rows of Dieter cells and other supporting cells (Fig. 1E). Finally, CHD4 was strongly expressed in the nucleus of SGNs marked by TUBB3 in the modiolus (Fig. 1F). Expression of CHD4 was present in all TUBB3-marked SGNs. The presence of *Chd4* transcripts in developing SGNs (Lu, Appler et al. 2011) and our results suggest that CHD4 may function in SGNs during inner ear development.

### *Chd4* is dispensable for initial SGN viability

CHD4 is present in many inner ear cell types, and mutations that inactivate CHD4 chromatin remodeling activity may contribute to improper development or dysfunction of multiple inner ear cell types. Since SGNs highly express CHD4, we sought to delineate how CHD4 ablation in SGNs contributes to cochlear development and hearing loss. Targeted ablation of CHD4 from SGNs using a *Ngn1* CreER^T2^ *Chd4*^flox/flox^ animal was accomplished. The *Ngn1* CreER^T2^ *Chd4*^flox/flox^ animals contain two copies of the *Chd4* cKO allele, where loxP sites flanked exons that code for the ATPase domain required for nucleosome repositioning (Williams, Naito et al. 2004). The *Ngn1* CreER^T2^ transgenes harbor a tamoxifen-inducible Cre expressed in neurosensory progenitors and can be used for genetic manipulations in SGNs (Koundakjian, Appler et al. 2007). Tamoxifen administration to *Ngn1* CreER^T2^ animals enabled Cre-mediated excision of loxP-flanked DNA in inner-ear neurosensory progenitors that give rise to SGNs (Koundakjian, Appler et al. 2007, Raft, Koundakjian et al. 2007). To ensure Cre activity was present in the intended cell types, *Chd4*^flox/flox^ *Ngn1* CreER^T2^ were crossed to the Ai9 tdTomato reporter (Madisen, Zwingman et al. 2010). In the triple-transgenic animals, tamoxifen administration induces Cre-activated Ai9 tdTomato reporter expression and enable fluorescence visualization of SGN cell bodies and axonal projections while simultaneously deleting *Chd4* (Fig. S1A). Control (*Ngn1* CreER^T2^; Ai9) and *Chd4* cKO (*Ngn1* CreER^T2^; *Chd4*^flox/flox^; Ai9) lines were generated. Timed matings with either the control or *Chd4* cKO lines produced pregnant dams, which were administered tamoxifen to produce embryos.

During inner ear development, delaminating otic progenitors around E9 from the proneurosensory domain of the otic vesicle coalesce to form the cochlear-vestibular ganglion (CVG). As development progresses, the CVG neurons segregate to create the spiral ganglion. A population of neurons exits the cell cycle and terminally differentiates at the base and middle of the cochlea, starting from E9.5-10.5. Most SGNs exit the cell cycle around E11.5 in the middle and base of the cochlea, while cells in the apex exit the cell cycle at E12.5. After exiting the cell cycle, peripheral projections from SGNs extend toward the sensory epithelium of the cochlea. At E12.5, SGN projections begin to extend beyond the spiral ganglion border. At E15.5, SGN peripheral axon outgrowth continues along the length of the cochlea. Type I SGNs form radial fiber bundles that undergo fasciculation. Between E15.5-18.5, type I-like SGN processes show extensive branching around and beyond the inner hair cells. The neurites are subsequently refined by retracting from the outer hair cell region (Appler and Goodrich 2011, Coate, Spita et al. 2015). Type II SGNs are notably distinct from type I SGNs by E16.5 based on their peripheral projections ending at the outer hair cell region (Bruce, Kingsley et al. 1997). The *Ngn1* CreER^T2^ animals with a reporter can mark delaminating neurosensory progenitors that develop into type I and type II SGNs at the aforementioned developmental stages (Koundakjian, Appler et al. 2007).

*Chd4* was deleted by administering daily tamoxifen to pregnant dams between E8.5 and E10.5, resulting in recombination in neurosensory progenitors that give rise to SGNs. Labeled cells were used to study the consequences of *Chd4* ablation in developing type I and type II SGNs. E18.5 embryos were harvested from pregnant dams after tamoxifen treatment. Although a subset of embryos survived to late embryonic stages, a substantial fraction underwent resorption prior to birth after tamoxifen administration. Quantification and comparison of resorbed embryos from control and *Chd4* cKO dams suggest a significant reduction in viability of *Chd4* cKO embryos compared to controls (Fig. S1B). Consistent with this, the number of surviving embryos from individual litters obtained from pregnant dams showed a statistically significant reduction in embryo viability after *Chd4* deletion (Fig. S1C). Cre recombinase activity in the *Ngn1* CreER^T2^ is not limited to the inner ear, but can also be observed throughout the central nervous system (Kim, Hori et al. 2011), suggesting that reduced viability likely reflects broader requirements for CHD4 during development.

Cochleae were harvested from the remaining embryos and used for immunostaining. To ensure that the genetic deletion of *Chd4* resulted in a lack of protein, immunofluorescence labeling for the CHD4 protein was performed. TUBB3 immunofluorescence was used to identify SGNs. The tdTomato fluorescence suggested that Cre activity was present in TUBB3-marked SGNs in both control and *Chd4* cKO cochleae samples. Even though SGNs were present in the cochleae, only the CHD4 signal was seen in controls and not in *Chd4* cKO cochleae (Fig. 2A). These results showed that CHD4 protein was absent in the *Chd4* cKO cochlea after tamoxifen treatment and suggest that CHD4 was not essential for SGN viability at embryonic and neonatal stages. In the E18.5 embryonic cochleae, control and *Chd4* cKO mice had similar numbers of tdTomato-marked SGNs (Fig. S2A). To quantify the percentage of the marked cell populations, SGNs from dissected cochleae were divided into three regions corresponding to the apex, middle, and base of the cochleae. In controls, a low percentage of cells lacking TUBB3 but expressing tdTomato was observed (apex: 6.81 ± 1.69%, middle: 4.07 ± 1.94%, base: 3.95 ± 1.74%), while in *Chd4* cKO cochlea, increased percentages of these cells were observed along the length of the cochleae (apex: 12.06 ± 2.25%, middle: 8.73 ± 0.91%, base: 7.82 ± 1.29%; apex, p < 0.14, middle, p < 0.10; base, p < 0.15) (Fig. S2B). The percentage of TUBB3-labeled SGNs marked by tdTomato confirmed the efficiency of Cre-mediated activity in these cells. In control embryos, Cre activity was observed in almost all SGNs with a slight decrease in labeled cells at the apex (apex: 89.86 ± 3.56%, middle: 97.09 ± 1.48%, base: 98.97 ± 0.36%). Since development proceeds in a base-to-apex manner, the decreased percentage of tdTomato-marked cells located at the apical region of the cochlea was likely due to a small population of late-born neurons that were not exposed to tamoxifen during the time window of administration. *Chd4* cKO displayed similar percentages of tdTomato-marked cells (apex: 84.35 ± 2.99%, middle: 95.71 ± 1.96%, base: 98.91 ± 0.2%; apex p < 0.30; middle p < 0.60; base, p < 0.90). The percentages of tdTomato+TUBB3+ cells from control and *Chd4* cKO cochleae were not statistically different (Fig. S2C). The number of tdTomato+ cells from *Chd4* cKO was not significantly different from control (44.22 ± 0.41% for control, 49.70 ± 2.32% for *Chd4* cKO, p < 0.08) (Fig. S2D). These results suggest that tamoxifen-induced Cre recombination was efficient along the entire length of the cochlea and was similar between control and *Chd4* cKO cochleae. Furthermore, *Chd*4 is not required for SGN survival.

**Figure 2.**
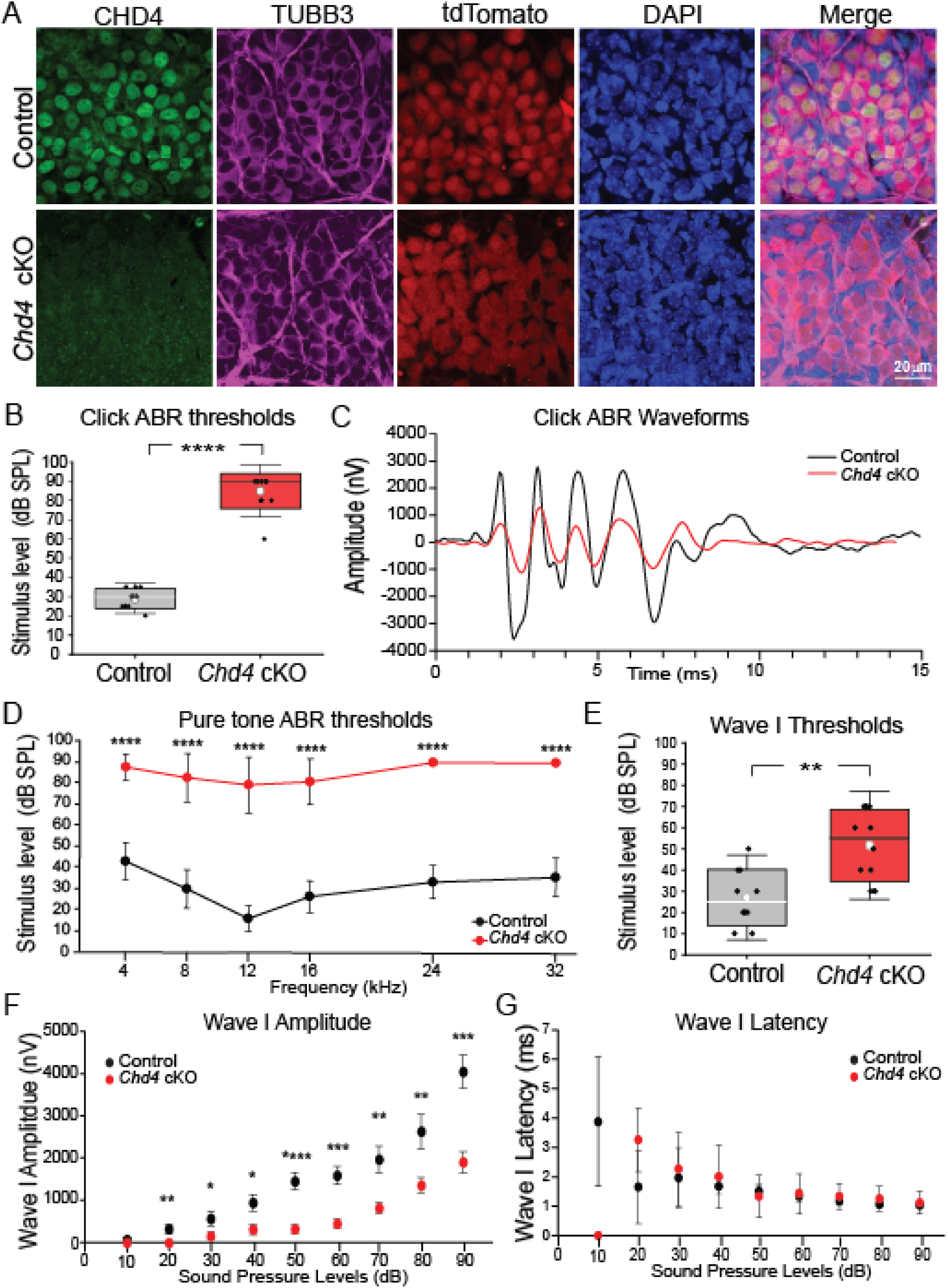
Auditory brainstem response (ABR) defects of *Chd4* cKO mice. (A) Immunostaining of CHD4 and TUBB3 marked neurons in the E18.5 cochlea. tdTomato expression correlates with the presence of Cre activity. Nuclear CHD4 expression was observed in DAPI-marked nuclei in control SGNs but was absent in *Chd4* cKO SGNs after tamoxifen administration. (B) Box-and-whisker plots show mean click ABR thresholds for control and *Chd4* cKO (p < 1.65 × 10^-13^). (C) Representative ABR waveforms of control and *Chd4* cKO mice in response to a click sound stimulation at 90 dB. (D) Pure-tone ABR thresholds measured at 4- 32 kHz. *Chd4* cKO mice exhibited significantly elevated thresholds across all tested frequencies (all p < 1 × 10^-10^). (E) Box-and-whisker plots show mean ABR Wave 1 threshold values for control and *Chd4* cKO (p < 0.01). (F) Wave 1 amplitudes measured across a range of click sound pressure levels. Amplitudes were significantly reduced in *Chd4* mice at all intensities above 20 dB SPL (10 dB, p <0.12; 20 dB, p <4.36 × 10^-3^; 30 dB, p < 0.04; 40 dB, p < 1.28 × 10^-2^; 50 dB, p < 5.82 × 10^-5^; 60 dB, p < 1.04 × 10^-4^; 70 dB, p < 1.92 × 10^-3^; 80 dB, p < 7.32 × 10^-3^; 90 dB, p < 1.21 × 10^-4^). (G) Wave 1 latencies were largely unchanged between genotypes (30 dB, p < 0.45; 40 dB, p < 0.11; 50 dB, p < 0.79; 60 dB, p < 0.79; 70 dB, p < 0.17; 80 dB, p < 0.15; 90 dB, p < 0.30). The Student’s *t*-test was used for statistical analysis with the number of animals (control, n=10; *Chd4* cKO, n=12). If the p-value is not reported, the statistical analysis could not be performed because no detectable values were observed in *Chd4* cKOs. Data are presented as mean ± standard deviation. Scale bar as marked.

### *Chd4* cKO animals showed altered auditory brainstem responses

We evaluated the physiological function of SGNs using auditory brainstem responses (ABRs). Wave I of the ABR corresponds to the synchronous firing of SGNs within the auditory nerve and serves as a readout. Given the widespread tonotopic distribution of tdTomato-positive SGNs, we used a broadband click stimulus to assess overall auditory function. Click-evoked ABRs measured across a sound pressure level (SPL) range of 10 to 90 dB revealed a profound increase in hearing thresholds in *Chd4* cKO mice compared to control littermates (29 ± 90.5 dB SPL for control and 85 ± 5.16 dB SPL for *Chd4* cKO, p < 1.65 × 10^-13^) (Fig. 2B), accompanied by a noticeable reduction in ABR waveform amplitudes (Fig. 2C). To assess frequency-specific deficits, we delivered pure-tone stimuli in octave steps between 4 and 32 kHz. Consistent with our click-evoked responses, *Chd4* cKO animals exhibited a significant increase in ABR thresholds across all frequencies (4- 32 kHz; all p <1 × 10^-10^), demonstrating an auditory deficit across the tonotopic axis (Fig. 2D). Given that the *Ngn1*-CreER^T2^ line targets *Chd4* ablation specifically to SGNs, we next analyzed Wave I amplitudes to evaluate SGN outputs. We observed a significant increase in Wave I thresholds in *Chd4* cKO animals relative to controls (27 ± 13.37 dB SPL for control and 51.67 ± 16.97 dB SPL for *Chd4* cKO, p < 1.65 × 10^-13^) (Fig. 2E). In agreement with increased thresholds, Wave I amplitudes were significantly reduced in *Chd4* cKO mice across nearly all sound intensities (Fig. 2F). At the lowest SPL (10 dB), Wave I peaks were not detectable in *Chd4* cKO animals. In contrast to the significant reduction in Wave I amplitudes, Wave I latencies did not differ between genotypes across most sound pressure levels, except at 10 dB SPL, where waveforms were not observed in *Chd4* cKO animals (Fig. 2G). Taken together, these data demonstrate that in SGNs, *Chd4* is required for SGN function.

### *Chd4* cKO cochleae displayed altered fasciculation of radial fiber bundles

To determine the cellular consequences of *Chd4* deletion, we evaluated SGN morphology to ascertain whether the observed functional attenuation stems from primary structural defects or a disrupted auditory circuit. SGN peripheral innervation was determined by immunofluorescent labeling to observe the initial part of the auditory circuit. Post-mitotic neurons from the spiral ganglion extend neurites towards inner and outer hair cells, starting from the base to the apex. Type I and type II SGNs target IHCs and OHCs, respectively. Type I and type II SGN fiber tracks and innervation patterns were marked by performing whole-mount TUBB3 immunostaining using cochleae from control and *Chd4* cKO (Fig. 3A).

**Figure 3.**
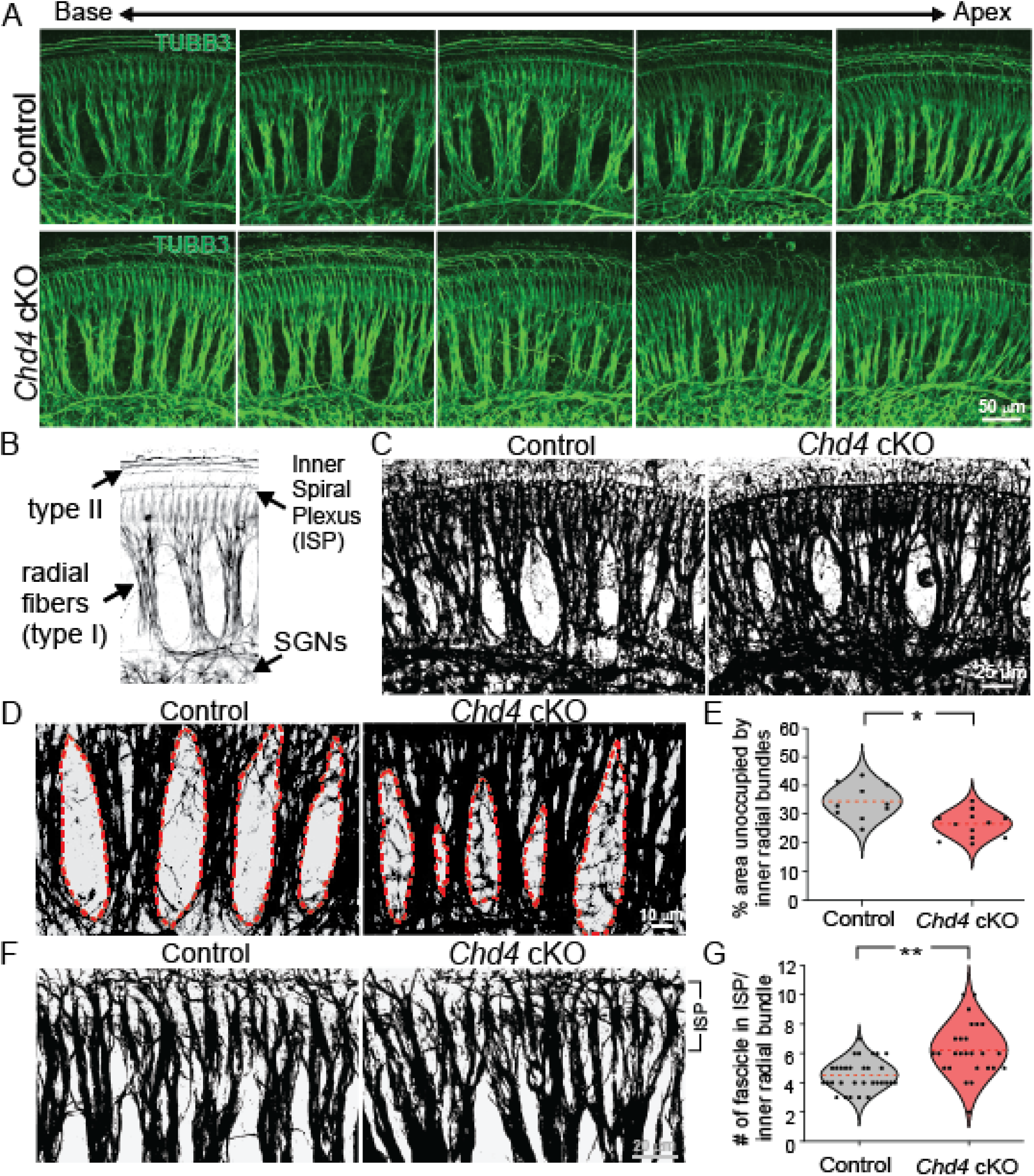
Inner radial bundles display fasciculation defects in *Chd4* cKO cochleae. (A) TUBB3-labeled cochleae from E18.5 control and *Chd4* cKO animals. Images are ordered from base to apex. (B) Depiction of the inner spiral plexus (ISP) near the base of inner hair cells and peripheral projections from type I (radial fibers) and II SGNs from E18.5 cochlea. (C) Binarized images of TUBB3-labeled control and *Chd4* cKO cochleae at E18.5. (D) Regions between radial fiber bundles were outlined in red. (E) The identified areas were quantified and compared between control and *Chd4* cKO (p < 0.05; control, n=10; Chd4 cKO, n=13 images were analyzed from control, n=3; and *Chd4* cKO, n=3 cochleae). (F) Binarized images of control and *Chd4* cKO E18.5 cochleae around the ISP. (G) Comparison of the normalized number of fascicles in the ISP per radial bundle between control and *Chd4* cKO cochleae (p < 0.01; control, n=31, and *Chd4* cKO, n=26 bundles were analyzed from control, n=5, and *Chd4* cKO, n=3 cochleae). The Student’s *t*-test was used for statistical analysis. Scale bars as marked.

The peripheral axons of type I SGNs extend from the soma to form a radial bundle. The type I SGN fibers fasciculate, the terminals extensively branch around the inner spiral plexus (ISP), and retract from the outer hair cell region back into the IHC region (Fig. 3B). In *Chd4* cKO cochleae, radial fibers were less compact and showed wider and less compact nerve fascicles. The area between the fiber bundles was identified by converting TUBB3 immunofluorescent images into binary black-and-white images. The white regions between fiber bundles and the total area from the SGN soma to the sensory epithelium were determined from images taken along the length of the cochlea (Fig. 3C). The percent area unoccupied by radial fibers relative to the total area was calculated and used as a metric for fasciculation (Fig. 3D). The average percentage of unoccupied space in *Chd4* cKO cochleae was significantly decreased (33.19 ± 2.59% for control and 25.12 ± 1.28% for *Chd4* cKO, p < 0.05). The result suggested that the radial fibers from *Chd4* cKO cochleae covered more area and thus were wider than controls (Fig. 3E). The fiber bundles were counted in control and *Chd4* cKO cochleae to determine the fascicle numbers in the inner spiral plexus (ISP) (Fig. 3F). The ISP consists of the region where radial fibers extend from the soma through the *habenula perforata* to innervate the IHC (Fig. 1). The average number of fascicles in ISP per inner radial bundle increased significantly in *Chd4* cKO cochleae (4.48 ± 0.16 bundles for control and 6.54 ± 0.67 bundles for *Chd4* cKO, p < 0.01) (Fig. 3G). These results show an increased number of inner radial fiber bundles and less compact bundling of the nerve fiber in the absence of *Chd4* in SGNs.

### *Chd4* cKO cochleae display improper turning of type II fibers

Type II SGNs constitute only 5-10% of the total SGN population, but their axonal projections are spatially distinct from type I SGNs. During development, their peripheral processes cross the tunnel of Corti into the outer hair cell region, where the axons make a right-angle turn and travel from the base of the cochlea towards the apex to make *en passant* synapses with outer hair cells. The outer spiral fibers are the bundled neurites from type II SGNs that extend along the OHC region. Each OHC is innervated by 2-5 type II SGNs (Huang, Barclay et al. 2012, Martinez-Monedero, Liu et al. 2016). To look at the outer spiral fiber tracks, TUBB3-labeled neurons from whole-mount cochleae were used for analysis (Fig. 4A). The outer spiral fibers usually display three main fiber tracks that travel along individual rows of outer hair cells with intermittent fibers crossing these tracks. To show quantitative differences between the outer spiral tracks, intensity profiles from control and *Chd4* cKO cochleae images spanning the outer hair cell region were taken (Fig. 4B). Measurements of the intensity profile show different peaks corresponding to the presence of fibers. The intensity profile plot showed three prominent peaks in the control cochlea (black). In contrast, *Chd4* cKO cochlea (red) displayed additional peaks (Fig. 4C). The data suggested that some type II SGN fibers or individual axons that detract from the outer fiber tracts (Fig. 4C). Each major peak corresponds to an outer spiral bundle consisting of multiple fibers. The appearance of the minor peaks within a major peak corresponds to the increased dispersion of the fibers within an outer spiral fiber. The number of distinct fibers observed by fluorescence microscopy in the outer hair cell region was counted and normalized to area to validate the altered axon paths. Similar to the intensity profile, the number of fibers in the OHC area increased in *Chd4* cKO samples compared to controls (13.98 ± 0.73 fibers for controls and 17.64 ± 0.78 fibers for *Chd4* cKO, p < 0.03) (Fig. 4D). These results suggest aberrant bundling of the outer spiral fibers after *Chd4* ablation in SGNs.

**Figure 4.**
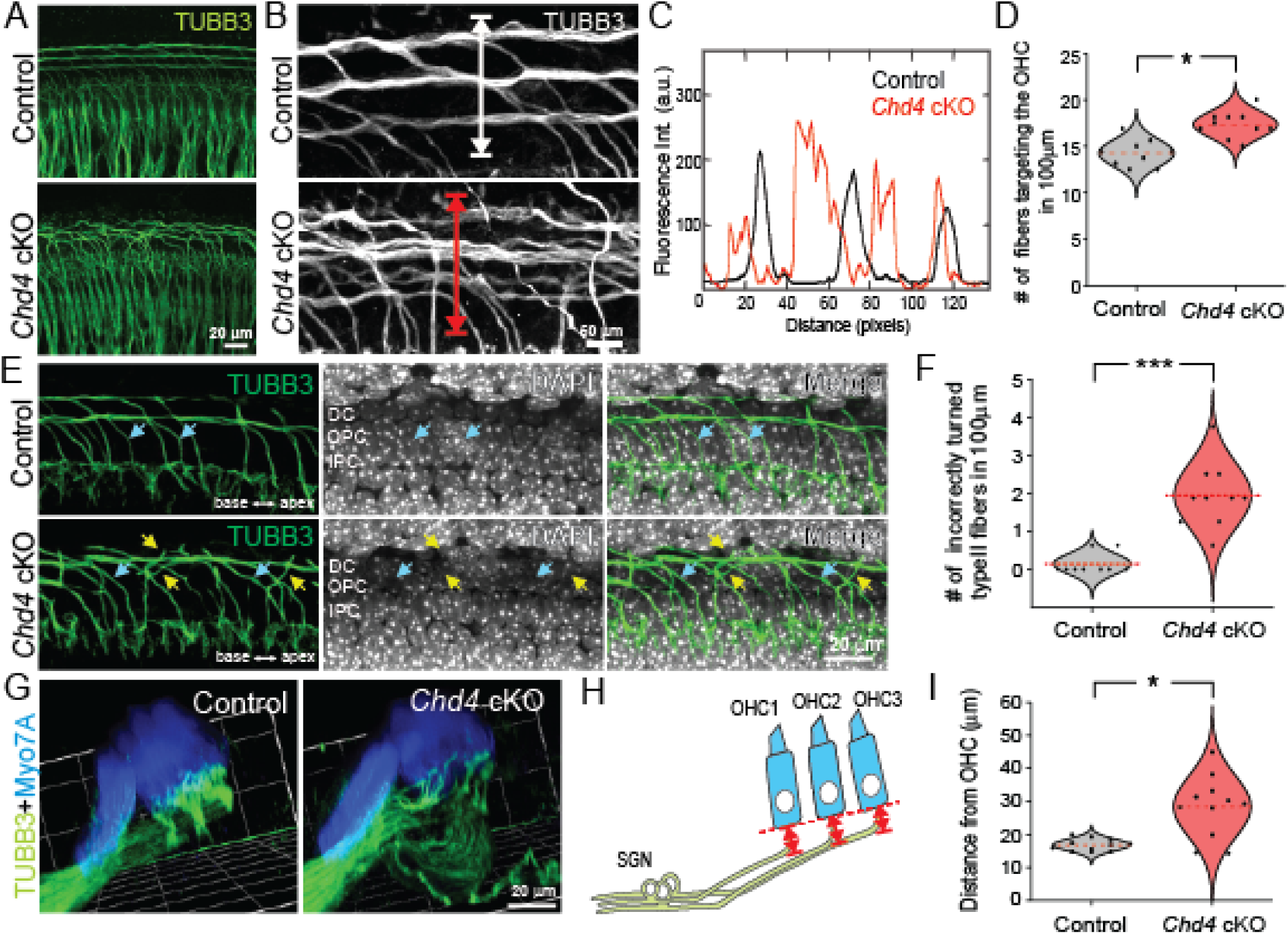
Type II spiral ganglion neurons display turning defects in the *Chd4* cKO mice. (A) TUBB3-labeled neuronal projections in E18.5 control and *Chd4* cKO cochleae. Type II fibers can be distinguished by their anatomical location. (B) Type II fibers from the control show stereotypic tracks that turn towards the base and travel along individual rows of outer hair cells. *Chd4* cKO type II fibers showed aberrant fiber tracks. (C) The intensity profile plot was measured along the white and red lines to highlight the differences between the control and *Chd4* cKO fiber tracks. Peaks from fluorescence intensity measurements indicate the presence of a fiber track. Control cochleae show three peaks correlating with fibers along the three rows of outer hair cells. *Chd4* cKO samples showed an additional peak, suggesting aberrant fiber tracks. (D) Normalized number of fibers entering the OHC region per 100 µm from control and *Chd4* cKO E18.5 cochleae (p < 0.03; control, n=9, and *Chd4* cKO, n=10 images were analyzed from control, n=3, and *Chd4* cKO, n=3 cochleae). (E) Optical section from TUBB3 and DAPI-labeled confocal micrographs that visualize fibers entering the OHC region. Blue arrows mark correctly turned, while yellow arrows show incorrect turning type II fibers. (F) The normalized number of incorrectly turned type II fibers per 100 µm from control and *Chd4* cKO E18.5 cochleae (p < 2.82 × 10^-4^; control, n=3, and *Chd4* cKO, n=3 cochleae). (G) 3D-rendered images highlighting the axonal trajectory from SGNs (TUBB3+, green) to hair cells (MYO7A+, blue). (H) Schematic of peripheral SGN axonal projections to outer hair cells. Red arrows represent the distance between contact points in the OHCs (red dashed line) and the final turning point on the incoming axon. (I) Average distances from the base of outer hair cells to the axon bundles showed differences between control and *Chd4* cKO (p < 0.03; control, n=11, and *Chd4* cKO, n=11 images were analyzed from control, n=3, and *Chd4* cKO, n=3 cochleae). The Student’s *t*-test was used for statistical analysis. Scale bars as marked.

We also noticed the aberrant turning of the outer spiral fiber tracks. Turning of the type II SGN fibers usually occurs near the inner pillar cell (IPC), outer pillar cell (OPC), or Dieter cell (DC) region. Confocal micrographs were acquired below the outer hair cell layer up to the sensory epithelium to visualize the fiber turning. Individual fibers from control cochleae turned towards the base of the cochlea before joining one of the three outer spiral fiber tracks. In contrast, the *Chd4* cKO cochleae showed fibers turning in the opposite direction (Fig. 4E, yellow arrows). Incorrectly turned type II fibers were quantified in control and *Chd4* cKO cochleae (0.12 ± 0.06 fibers for control and 1.93 ± 0.14 fibers for *Chd4* cKO, p < 2.83 x 10^-4^) (Fig. 4F). These data showed that the deletion of *Chd4* increases the incorrect turning of type II SGNs in the outer spiral fibers. Turning of the type II fibers has previously been shown to require signals from supporting cells (Ghimire, Ratzan et al. 2018, Ghimire and Deans 2019, Deans 2022). To ensure that supporting cells in the organ of Corti were present and not indirectly affected by *Chd4* deletion, immunostaining for both hair cells and supporting cells, using MYO7A and SOX2, was performed in control and *Chd4* cKO cochleae. A single row of inner hair cells and three rows of outer hair cells were present in both samples. Moreover, SOX2-labeled supporting cells showed the presence of organized supporting cells residing below hair cells (Fig. S3). These results suggest that *Chd4* deletion in SGNs affects the turning of type II SGNs in a cell-autonomous manner.

### *Chd4* cKO cochleae display aberrant basal to apical axon path towards outer hair cells

Alterations in axon paths could also be due to how fibers exit Rosenthal’s canal and ascend toward the outer hair cells. To determine if this is the case, reconstructed confocal images of MYO7A-labeled hair cells provide a landmark for the terminal destination of axons. We observed that many type II tracks take a more circuitous route toward the OHCs because the fibers initially descend basally and migrate further from the OHC soma (Fig. 4G). The distances were obtained from the bottom of individual OHC somas to the furthest detectable axon fibers directly below. The measurements were used to quantify the aberrant axon path (Fig. 4H). The measurements showed an increase in the average distance of the fibers to the bottom of hair cells in *Chd4* cKO cochleae (17.84 ± 0.66 µm for control and 29.87 ± 3.74 µm for *Chd4* cKO, p < 0.03). The increased distance measured in *Chd4* cKO cochleae suggested that outer spiral fibers do not properly ascend from Rosenthal’s canal towards the outer hair cells (Fig. 4I). The fiber tract and innervation changes observed in *Chd4*-deficient type I and type II SGNs implicate CHD4 in axon pathfinding. However, its molecular targets in SGNs have not yet been identified.

### Enrichment of CHD4 at cis-regulatory regions of *Eph* and *Ephrin* genes

CHD4 function is cell-type dependent, and its chromatin activity likely alters a unique repertoire of cis-regulatory elements in SGNs to control gene expression. A significant hurdle is identifying high- confidence CHD4 occupancy sites in SGNs using chromatin immunoprecipitation followed by deep sequencing (ChIP-seq) or Cleavage Under Targets & Tagmentation (CUT&Tag). The inability to identify these high-confidence sites in SGNs is partly due to the large number of cells required for the experiments. Instead of primary SGNs, we used immortalized multipotent otic progenitor (iMOP)-derived neurons to map CHD4 genome-wide binding sites and identify potential target genes. Immortalized multipotent otic progenitors (iMOP) were derived from primary cochlear otic progenitors and have highly similar transcriptomes. iMOPs can differentiate into cells that display bipolar and pseudounipolar morphology similar to SGNs (Kwan, Shen et al. 2015).

Immunostaining confirmed the presence of CHD4 in the nucleus of proliferating iMOPs and TUBB3-expressing iMOP-derived neurons (Fig. 5A). To identify genome-wide CHD4 binding sites, CUT&Tag was performed using CHD4 antibodies on proliferating iMOP and iMOP-derived neurons. CHD4 occupancy is cell type-specific and is recruited to active promoters and enhancers to repress transcription (Yang, Yamada et al. 2016). We used the H3K4me3 and p300 marks as genomic landmarks for promoters and enhancers. Sparse Enrichment Analysis for CUT&RUN (SEACR) identified high-confidence enrichment regions, known as peaks, across all CUT&Tag sequence reads (Meers, Bryson et al. 2019). We compared CHD4 peaks to H3K4me3 and p300-bound regions. The heatmaps show the distribution of reads within a ±3 kb window. The profile plots allowed us to qualitatively evaluate read density across the same genomic regions. We identified CHD4+ H3K4me3+ regions based on the occupancy of CHD4 and H3K4me3 peaks in proliferating iMOP cells and iMOP-derived neurons (Fig. S4B). The CHD4+ H3K4me3+ regions corresponded to CHD4-bound promoters. Similarly, we defined CHD4+ p300+ regions in proliferating iMOP cells and iMOP-derived neurons (Fig. S4C). The CHD4+ p300+ regions corresponded to CHD4-bound enhancers. From the profile plots, we observed that the CHD4 signal was higher in iMOP-derived neurons than in proliferating iMOPs at both promoters and enhancers. We quantified and compared the CHD4 total signals at CHD4+ H3K4me3+ promoters and CHD4+ p300+ enhancers between proliferating iMOP and iMOP-derived neurons. We showed that at CHD4-bound promoters, CHD4 total signals increased 2.80-fold in iMOP-derived neurons (57,811.40 ± 65,983.40 RPKM) compared to proliferating iMOPs (20,684.90 ± 25,408.89 RPKM, p < 2.20 x 10^-16^). At CHD4-bound enhancers, we found a 4.65-fold increase of CHD4 total signal in iMOP-derived neurons (134,027.50 ± 102,349.97 RPKM) compared to proliferating iMOPs (28,837.65 ± 29,014.11 RPKM, p < 2.20 × 10^-16^) (Fig. 5B). The results suggest that increased enrichment of CHD4 at cis-regulatory regions in iMOP-derived neurons may regulate gene expression during neuronal differentiation.

**Figure 5.**
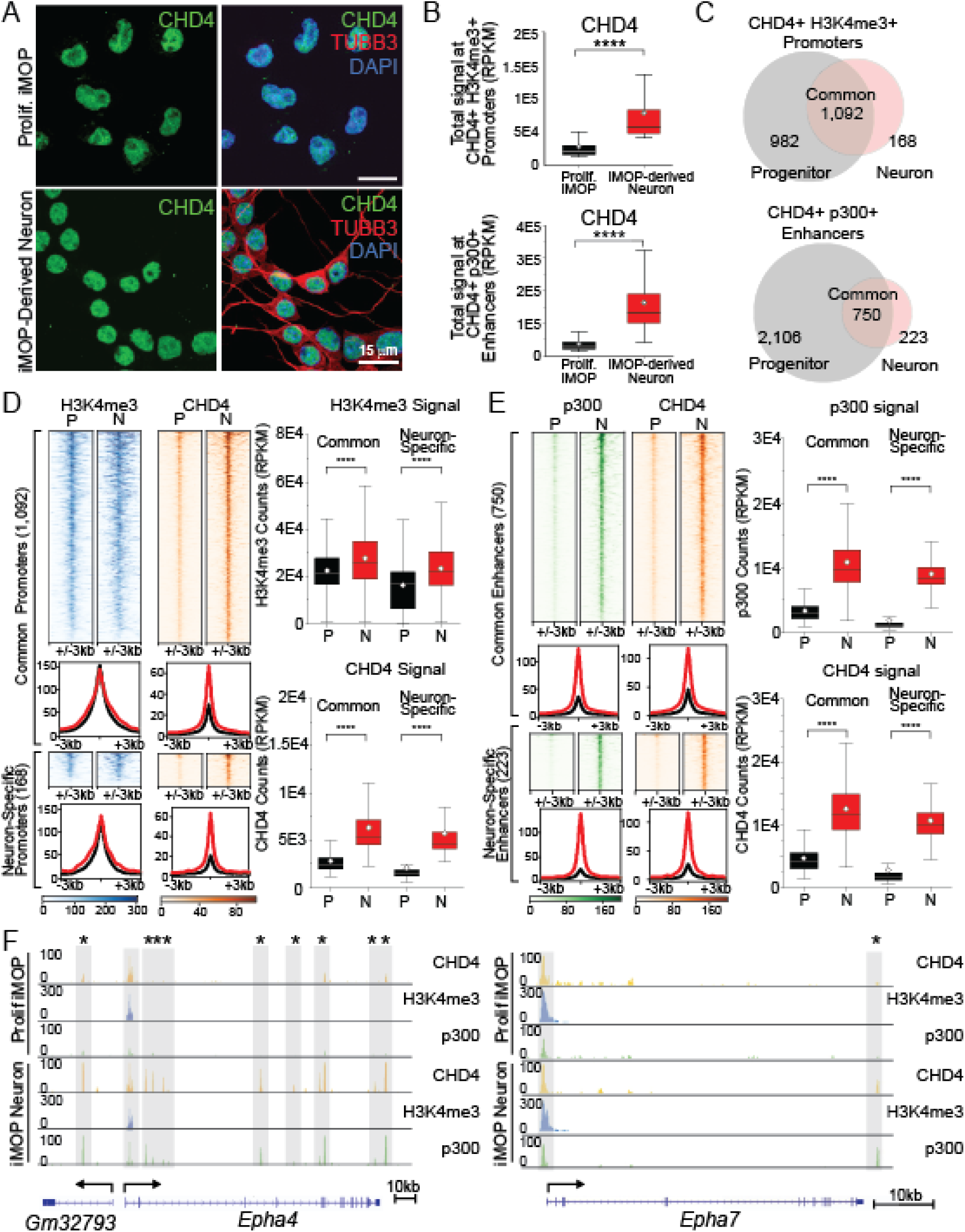
CHD4 binds to the promoters of genes involved in axon guidance, including members of the ephrin family of signaling molecules, in iMOP-derived neurons. (A) Overlap of CHD4 in the nuclei (DAPI) of proliferating iMOP and TUBB3-labeled iMOP-derived neurons. CUT&Tag was performed using proliferating iMOP cells and iMOP-derived neurons. CUT&Tag peaks were identified using SEACR. (B) Box-and-whisker plots show that SEACR total signals for CHD4 at CHD4+ H3K4me3+ and CHD4+p300+ in proliferating iMOPs were significantly different from those in iMOP-derived neurons (p < 2.20 × 10^-16^). (C) Venn diagrams show the overlap of CHD4+ H3K4me3+ promoters and CHD4+ p300+ enhancers between progenitors (P) and neurons (N). (D) Heatmaps and profile plots show H3K4me3 and CHD4 signals within ± 3kb of the summit regions of common and neuron-specific promoters. Box-and-whisker plots show significantly increased H3K4me3 and CHD4 signals at both common and neuron-specific promoters in progenitors compared to neurons. (E) Heatmaps and profile plots show p300 and CHD4 signals within ± 3kb of the summit regions of common and neuron-specific enhancers. Box-and-whisker plots show significantly increased p300 and CHD4 signals at both common and neuron-specific enhancers in progenitors compared to those in neurons. (F) Enrichment of CHD4 (orange), H3K4me3 (blue), and p300 (green) at the specified genomic region of *Epha4* and *Epha7* in proliferating iMOPs and iMOP-derived neurons. Highlighted genomic regions represent H3K4me3-marked promoters. Highlighted regions with asterisks denote the regions with increased enrichment of CHD4 and p300 in iMOP-derived neurons. The number of asterisks denotes the peaks with increased CHD4 and p300 levels. The Wilcoxon rank-sum test was used for statistical analysis. Scale bars as marked.

To investigate the involvement of CHD4 in neuronal differentiation, we identified CHD4-binding sites by comparing CHD4-bound regions between proliferating iMOPs and iMOP-derived neurons. At CHD4+ H3K4me3+ promoters, we found 982 progenitor-specific, 168 neuron-specific, and 1,092 common promoters in both proliferating iMOPs and iMOP-derived neurons. At CHD4+ p300+ enhancers, we identified 2,106 progenitor-specific, 223 neuron-specific, and 750 common enhancers (Fig. 5C). We focused on common and neuron-specific regions since CHD4 activity may exert epigenetic changes at these sites during neuronal differentiation. We generated heatmaps and profile plots for each of the regions. At common promoters, we observed a 1.20-fold increase of H3K4me3 in iMOP-derived neurons (25,871.92 ± 12,797.10 RPKM) compared to proliferating iMOPs (21,550.36 ± 9,015.10 RPKM, p < 3.67 x 10^-23^) and a 2.28-fold increase in CHD4 in iMOP-derived neurons (5,407.49 ± 2,955.04 RPKM) compared to proliferating iMOPs (2,371.49 ± 1573.27 RPKM, p < 3.46 x 10^-282^). At neuron-specific promoters, a 1.31-fold increase in H3K4me3 (proliferating iMOP 17,105.85 ± 10,709.21 RPKM, iMOP-derived neurons 22,411.04 ± 14,180.57 RPKM, p < 3.87 x 10^-7^) and 3.30-fold increase in CHD4 (proliferating iMOP 1,397.09 ± 2,617.01 RPKM, iMOP-derived neurons 4,608.37 ± 3,173.27 RPKM, p < 4.22 x 10^-45^) were observed in iMOP-derived neurons (Fig. 5D). All observed increases in H3K4me3 and CHD4 were statistically significant.

We performed a similar analysis for enhancers. The heatmaps and profile plots showed distributions of p300 and CHD4 centered on common and neuron-specific enhancers. A 3.37-fold increase in p300 (proliferating iMOP = 2,838.41 ± 1,713.93 RPKM, iMOP-derived neuron = 9,578.39 ± 4531.32 RPKM, p < 5.92 x 10^-223^) was observed at common enhancers. CHD4 displayed a 2.84-fold increase in signal from iMOP-derived neurons (proliferating iMOP 4,068.79 ± 2,542.82 RPKM, iMOP-derived neuron 11,559.34 ± 4,647.43 RPKM, p < 9.30 x 10^-210^). At common and neuron-specific enhancers, a statistically significant increase in signal was detected for p300 and CHD4 in iMOP-derived neurons compared to proliferating iMOPs (Fig. 5E). The results suggest that CHD4 was significantly enriched at specific promoters and enhancers during neuronal differentiation. The increased CHD4 occupancy at promoters and enhancers likely alters the chromatin state of the cis-regulatory regions to regulate transcription.

To gain biological insights into CHD4 targets, genes near common and neuron-specific regulatory elements were identified (Table S1). The identified target genes were used for gene ontology analysis. Gene ontology (GO) analysis revealed a function in chromatin organization (p < 5.10 × 10-14), consistent with the predicted cellular process of CHD4 in inner ear neurons. Other biological processes such as ephrin receptor signaling (p < 4.90 x 10^-5^), axon guidance (p < 2.00 x 10^-13^), semaphorin-plexin signaling during axon guidance (p < 2.70 x 10^-2^), axonal fasciculation (p < 1.60 x 10^-2^), and axonogenesis (p < 1.60 x 10^-14^) were some of the notable cellular processes (Fig. S4A). The GO analysis suggested that CHD4 target genes may regulate processes involved in axon pathfinding, including ephrin receptor signaling. We identified *Epha4*, *Epha7*, *Ephb3*, *Efna3*, *Efna4*, *Efnb2*, *Efna5*, *Efnb1*, *Ephb2,* and other axon guidance genes as candidates based on their proximity to CHD4-bound promoter and enhancer regions (Table S1). Ephrin signaling pathways are essential in guiding SGN peripheral axons. We focused on *Epha4*, *Epha7*, and *Efnb2* because these genes were previously implicated in inner ear function (Coate, Raft et al. 2012, Defourny, Poirrier et al. 2013, Kim, Ibrahim et al. 2016, Petitpre, Wu et al. 2018, Sanders and Kelley 2022). At *Epha4* and *Epha7*, CHD4, and H3K4me3 peaks were observed near the transcriptional start site. We also observed an increase in CHD4 and p300 in iMOP-derived neurons at multiple intronic regions, a region upstream at *Epha4*, and a singular downstream site at *Epha7* (Fig. 5F). A similar observation was noted for *Efnb2* (Fig. S4D), where CHD4 was enriched at the H3K4me3-marked promoter along with increased CHD4 and p300 signals at a downstream enhancer region. In contrast, CHD4 was not observed at the transcriptional start site of *Calb2*, a non-target gene (Fig. S4E). These results implicate CHD4 in controlling the transcription of *Eph* receptors and *Efn* ligands. The altered expression of these genes may be pertinent to the axon-guidance deficits observed in the *Chd4* cKO cochleae.

### DNA binding motifs near Eph receptor and ephrin ligand genes

Although CHD4 may regulate the epigenetic state at cis-regulatory regions of axon guidance genes through its histone association domain, it lacks a sequence-specific DNA-binding motif (Laureano, Kim et al. 2023). We wanted to identify potential transcription factors that may regulate the expression of *Eph* and *Efn* genes. To gain insight into transcriptional regulation, *de novo* DNA motif discovery was performed on CHD4-enriched regions to identify potential transcription (TF) binding sites. Starting with SEACR-defined summits obtained from CHD4 peaks, we first identified 1,095 sites within ±1 kb of TSS (Fig. S5A). Next, we focused on CHD4 binding within ±1 kb of TSS at axon guidance genes to identify 16 sites (Fig. S5B). *De novo* motif discovery identified several motif consensus sequences. The motifs were annotated by comparing to known TF motifs, and statistically significant matches were displayed in a heatmap with their corresponding p-values (Fig. S5C). Motif occurrences and Information Content Scores (ICS), a conservation metric, were determined and plotted (Fig. S5D). Motif occurrences and relative positions to the TSS were shown for each axon guidance gene (Fig. S6A). The probability of base pair composition from identified motifs was visualized using sequence logos (Fig. S6B). Notably, motif occurrences were also identified in additional axon guidance genes, including members of the semaphorin (*Sema3a*, *Sema6c*, and *Sema7a*), plexin (*Plxna1* and *Plxna2*), and frizzled (*Fzd2*, *Fzd7*, and *Fzd8*) families (Fig. S6A).

We identified distinct combinations of TF motifs for *Epha4*, *Epha7*, and *Efnb2*, genes previously implicated in cochlear axon guidance. *Epha4* contained Stat2 (Signal transducer and activator of transcription 2), two Lhx3 (LIM homeobox 3), and an Ar (Androgen Receptor) motif. *Epha7* harbored Stat2, Plagl1 (Pleiomorphic Adenoma Gene-like 1), and Hnf1A (Hepatic Nuclear Factor 1 Alpha) motifs. *Efnb2* displayed a single *Ar* motif. These motifs were displayed as sequence logos (Fig. 6A). The analysis revealed candidate TFs that may coordinate with CHD4 to regulate transcription of *Epha4*, *Epha7*, and *Efnb2*. Together, the data suggest that CHD4 regulates chromatin accessibility at transcription factor-binding sites, modulating the expression of Eph receptors and Efn ligands to facilitate axon guidance.

**Figure 6.**
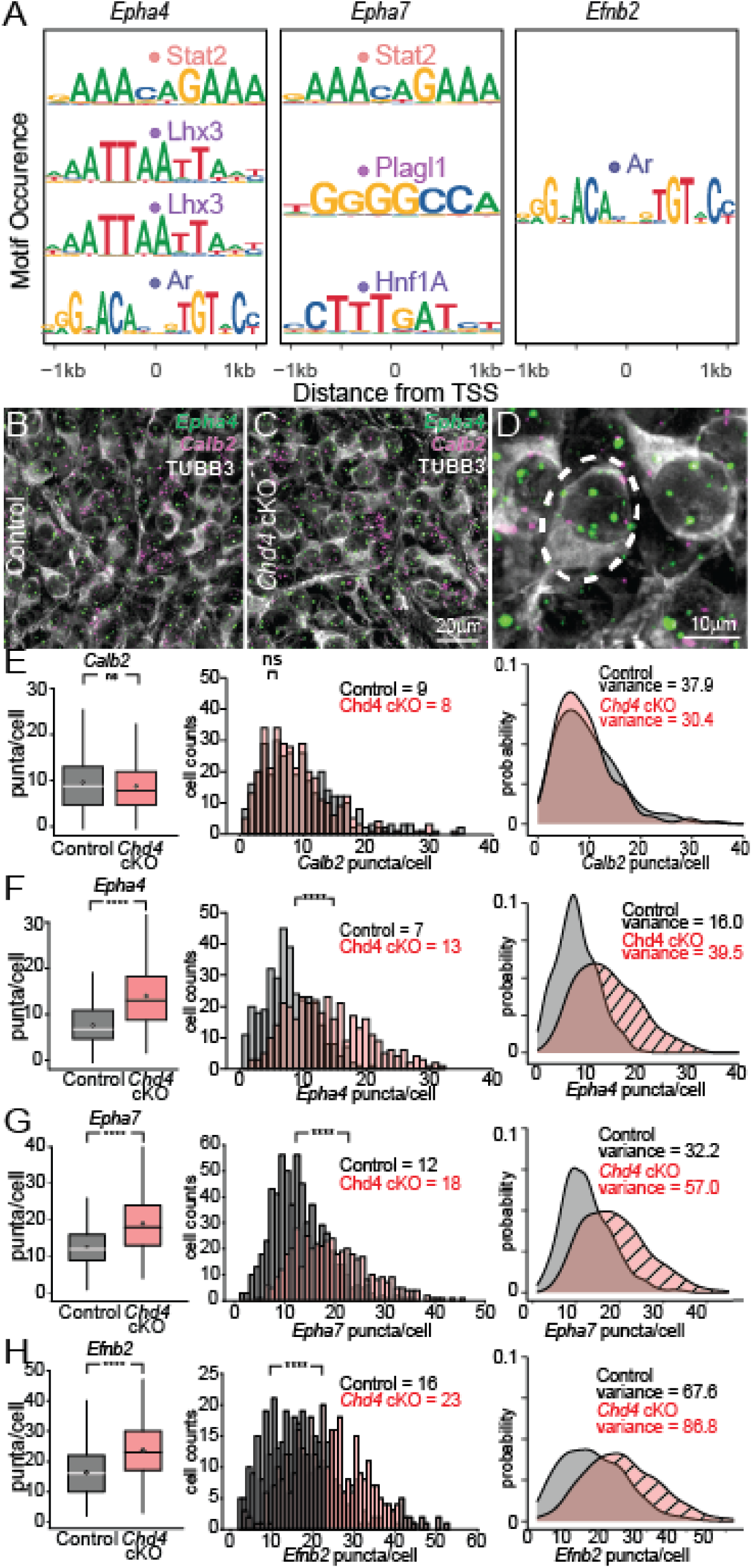
*Chd4* cKO shows an increased *Eph and Efn* mRNA expression in SGNs. (A) Occurrence and position of transcription factor (TF) binding motifs relative to the transcriptional start site (TSS) at *Epha4*, *Epha7,* and *Efnb2*. E16.5 (B) control and (C) *Chd4* cKO cochlea were processed for smFISH with probes against *Epha4* and *Calb2.* Cochleae were immunolabeled with TUBB3 to visualize SGNs. (D) The number of puncta for individual TUBB3-labeled cells (dashed line) was quantified. (E) Box plots representing median (lines) and mean (circles) values after comparing smFISH puncta between control and *Chd4* cKO samples for *Calb2* and distribution plots of puncta in control and *Chd4* cKO SGNs (p < 0.13). Kernel densities for *Calb2* transcripts (p < 0.04; control, n=350 cells from 3 cochleae and *Chd4* cKO, n=345 cells from 3 cochleae). (F) Box and distribution plots of the number of puncta for *Epha4* in control and *Chd4* cKO SGNs (p < 1.74 × 10^-39^). Kernel density plots for *Epha4* transcripts display the non-overlapping region (hashed region), suggesting an increased probability of cells from *Chd4* cKO with higher levels of *Epha4* transcripts compared to the control (p < 2.27 x 10^-14^; control, n=350 cells from 3 cochleae and *Chd4* cKO, n=345 cells from 3 cochleae). (G) Box and distribution plots of *Epha7* transcripts in control and *Chd4* cKO SGNs (p < 2.48 × 10^-45^). Non-overlapping region of densities in *Epha7* (hashed region) between control and *Chd4* cKO displays an increased probability of cells from *Chd4* cKO with higher levels of *Epha7* (p < 8.29 × 10^-11^; control, n=632 cells from 4 cochleae and *Chd4* cKO, n=436 cells from 3 cochleae). (H) Box and distribution plots of the number of *Efnb2* transcripts in SGNs from control and *Chd4* cKO (p < 7.33 × 10^-24^). Kernel density plots for *Efnb2* transcripts show an increased non-overlapping density region of *Efnb2* transcripts from *Chd4* cKO (hashed region) compared to the control. (p < 0.04; control, n=363 from 3 cochleae and *Chd4* cKO n=340 cells from 3 cochleae). The Wilcoxon rank-sum test and Levene’s test were used for statistical analysis.

### *Chd4* cKO showed increased *Eph* and *Efn* mRNA levels in SGNs

To determine whether deletion of CHD4 in SGNs affects ephrin levels, cochleae from control and *Chd4* cKO were subjected to single-molecule fluorescence *in situ* hybridization (smFISH). We used *Calb2* (*Calbindin2*) transcripts as a marker for SGNs and as a gene unaffected by CHD4. We simultaneously probed for the *Eph* or *Efn* transcripts. TUBB3 immunolabeling was combined with RNAscope to demarcate SGNs for quantification in control (Fig. 6B) and *Chd4* cKO (Fig. 6C). The soma of individual TUBB3-marked SGNs were used to determine regions of interest for puncta quantification to determine the relative number of mRNA molecules in each neuron (Fig. 6D).

The *Calb2* puncta were quantified, and there was no significant difference observed in *Calb2* transcript levels between *Chd4* cKO (8 ± 5.51 *Calb2* puncta/cell) and control (9 ± 6.16 *Calb2* puncta/cell, p < 0.13) (Fig. 6E). The distribution of *Calb2* puncta counts per cell was visualized as a histogram to display the number of cells with specified *Calb2* puncta/cell. This histogram was transformed into a density plot to illustrate the probability distribution of puncta counts per cell. The variance was calculated using the density plot. As observed, the density plots for both control and *Chd4* cKO SGNs were not appreciably different, and the variance of *Calb2* transcripts showed a slight 1.25-fold decrease in *Chd4* cKO (σ^2^ = 30.36) compared to the control (σ^2^ = 37.93, p < 0.04) (Fig. 6E).

In contrast, the average number of *Epha4* puncta in the SGN population increased in *Chd4* cKO (13 ± 6.29 puncta/cell) compared to the control (7 ± 4.00 puncta/cell, p < 1.74 × 10^-39^) and was statistically significant. By transforming the distribution of the *Epha4* puncta/cell into a density plot, we observed a 2.47-fold increase in the variance of *Epha4* transcripts in *Chd4* cKO (σ^2^ = 39.51) compared to control SGNs (σ^2^ = 16.00, p < 2.27 × 10^-14^). The non-overlapping regions in the density estimates reflect the increased probability of cells with higher *Epha4* transcript levels in the absence of *Chd4* (Fig. 6F). The results suggest that deletion of *Chd4* in SGNs increases *Epha4* in a subpopulation of SGNs. In contrast to *Epha4*, the decreased variance in *Calb2* puncta/cell may be caused by a CHD4-independent mechanism or an indirect effect of *Chd4* deletion.

To further validate additional axon guidance gene targets, we performed RNAscope on the two other members of the Ephrin receptor and ephrin ligand family, *Epha7* and *Efnb2*, which displayed CHD4 binding at their promoters. *Epha7* puncta in SGNs were significantly increased in *Chd4* cKO (18 ± 7.55 *Epha7* puncta/cell) compared to the control (12 ± 5.67 *Epha7* puncta/cell, p < 2.48 x 10^-45^) and the variance of *Epha7* transcripts showed a 1.77-fold increase in *Chd4* cKO (σ^2^ = 56.95) compared to the control SGNs (σ^2^ =32.19, p < 8.29 x 10^-11^) (Fig. 6G). A similar increase in *Efnb2* puncta *was* observed in *Chd4* cKO (23 ± 9.31 puncta/cell) compared to control (16 ± 8.22 puncta/cell, p < 7.33 x 10^-24^). A 1.28-fold increase in the *Efnb2* variance was observed in *Chd4* cKO (σ^2^ = 86.76) compared to control SGNs (σ^2^ =67.57, p < 0.04) (Fig. 6H). The results suggest an increased probability of *Chd4* cKO SGNs harboring more *Epha4*, *Epha7*, and *Efnb2* transcripts. The data indicate that CHD4 normally represses transcription of a subset of *Eph* receptor and *ephrin* ligand genes. Ablation of *Chd4* increases transcript levels in a subset of axon guidance genes in a proportion of cells, resulting in aberrant peripheral axon pathfinding.

## Discussion

CHD4 is expressed in cochlear supporting cells and hair cells from E18 to P21 (Layman, Sauceda et al. 2013). Gene expression analysis showed that *Chd4* transcripts are present and change in developing murine SGNs from E12 to P15 (Lu, Appler et al. 2011). *De novo* pathogenic variants of *CHD4* have been associated with SIHIWES, an autosomal dominant neurodevelopmental disorder. Patients with pathogenic variants of CHD4 show variable symptoms, but many have hearing loss (Sifrim, Hitz et al. 2016, Weiss, Terhal et al. 2016). CHD4 has a core ATPase SWI/SNF domain that hydrolyzes ATP and mediates chromatin repositioning. Some pathogenic mutations reduce the chromatin remodeling activity of CHD4 (Weiss, Lazar et al. 2020). To clarify the contribution of CHD4 in SGNs, we employed a cKO mouse model that enables Cre-dependent, inducible deletion of the exons encoding the ATPase domain of CHD4 in SGNs. The cKO mouse allowed us to separate CHD4 function from that of other inner ear cell types, such as hair cells and supporting cells. Our results implicate a cell-autonomous role for CHD4 in SGN peripheral axon pathfinding during inner ear development and highlight the importance of CHD4 function during embryonic development, when transcriptional programs governing axon guidance and neuronal wiring. The absence of CHD4 chromatin remodeling activity in SGNs led to hearing loss due to aberrant axon guidance. The altered pathfinding affects neural signal transmission and hair cell innervation, as evidenced by aberrant ABR responses. These findings link epigenetic changes to neuronal wiring and circuit formation.

CHD4 is a core subunit in the Nucleosome Remodeling and Deacetylase (NuRD) complex (Tong, Hassig et al. 1998, Xue, Wong et al. 1998, Zhang, LeRoy et al. 1998). The NuRD complex acts as a transcriptional repressor and localizes to sites of active transcription. At these sites, CHD4 repositions nucleosomes and subunits of the complex, such as histone deacetylases (HDAC) 1 and 2, which deacetylate histones (Morra, Lee et al. 2012, Watson, Mahajan et al. 2012). Although CHD4 has many other paralogs, distinct NuRD complexes with individual CHD paralogs (CHD3, CHD4, and CHD5) have specific molecular functions and do not compensate for the loss of a singular CHD. The remaining paralogous CHDs did not compensate for the cellular effects of ablating CHD4 (Nitarska, Smith et al. 2016). CHD4-containing NuRD complexes in the inner ear may contain distinct protein subunits that alter the epigenetic landscape at cis-regulatory regions. Our results suggest a role for CHD4 in axon guidance during SGN development. Even if paralogs were present in SGNs, they did not fully compensate for the observed axon guidance deficits.

Using iMOP-derived neurons as a cellular system for SGN development, we observed increased CHD4 occupancy at distinct sets of promoters and distal enhancers during neuronal differentiation. These findings suggest that CHD4 exerts epigenetic changes on cis-regulatory elements to attenuate transcription of axon guidance genes. Without CHD4, the core NuRD complex does not form, and axon guidance genes may show increased transcription without NuRD-dependent nucleosome repositioning and histone deacetylase activities. The exquisite temporal transcriptional control of axon guidance genes is likely required to establish the appropriate fasciculation and innervation patterns. Although our analysis focused on Eph/ephrin signaling, we observed CHD4 occupancy at multiple loci associated with axon guidance pathways. These include candidates that may participate in additional signaling systems, such as semaphorin pathways. While increased transcript levels were directly tested for selected Eph/ephrin genes, CHD4 binding was detected at loci associated with additional axon guidance pathways, suggesting a broader regulatory role. Coordinated regulation across multiple axon guidance pathways may explain how relatively modest transcriptional changes can lead to measurable axon guidance defects and severe physiological consequences, such as hearing loss.

CHD4 has domains that interact with modified histones, but it lacks a sequence-specific DNA-binding domain (Reid, Zhong et al. 2024). Motif analysis was performed to identify potential transcription factors (TFs) that may function with CHD4 at cis-regulatory regions. Motif analysis suggests candidate transcription factors may function with CHD4 at axon guidance gene loci. We identified TF binding motifs within a single CHD4 binding site at promoters of many axon guidance genes. The clustered motifs in some axon guidance genes, such as *Epha4* and *Epha7,* including transcription factor motifs such as *Lhx3*, *Stat2*, and *Hnf1A*, may facilitate cooperative TF binding. Individual transcription factor binding sites often exhibit relatively weak binding affinities. However, when multiple TFs bind nearby, they can engage in protein-protein interactions or induce DNA conformational changes that collectively strengthen their DNA-binding affinity. This cooperativity often leads to synergistic activation or repression of gene expression, in which the combined effect of multiple TFs exceeds the sum of each TF’s contribution (Wang, Xie et al. 2025). The synergistic binding of TFs with the nucleosome remodeling activity of CHD4 enables high specificity and precisely tunes transcriptional regulation of axon guidance genes. The distinct motifs within a cluster also highlight combinatorial control of transcription. These regulatory modules may serve as integration points for diverse cellular signals, allowing transcription only when a specific combination of TFs is present during development. This combinatorial logic could enable complex regulatory programs to coordinate distinct steps of axon guidance in the cochlea.

Our identified candidate set of Eph receptors and ephrin ligand genes (*Epha4*, *Epha7*, and *Efnb2*) is noteworthy, as it suggests that a combination of axon guidance genes contributes to the wiring of the developing cochlea. Eph receptors are a family of receptor tyrosine kinases that mediate many essential processes through promiscuous interactions with membrane-bound ephrin ligands (Kania and Klein 2016). Identifying Eph receptors and their ephrin ligands in the cochlea is particularly challenging. The cochlea’s intricate structure makes isolating specific Eph/ephrin interactions complicated without disrupting the function of other cell types in the tissue. The Eph/ephrin system also features significant redundancy and promiscuity, where multiple receptors can bind to the same ligand and vice versa, complicating the identification of specific functional roles. Genetic knockout models often reveal compensatory mechanisms among Eph family members, masking phenotypic effects and complicating functional studies. Here, we used epigenetic modifications of regulatory elements surrounding a candidate set of axon guidance genes that are responsible for assembling neural circuits in the cochlea. Ephrins are divided into the glycosyl-phosphatidylinositol (GPI) linked ephrin-A and the transmembrane ephrin-B ligands. Ephrins (EFN) bind to Eph receptors (EPH) of the same class, except for EPHA4. EPHA4 interacts with the EFNA B2 and B3 ligands. Deletion of *Epha4* showed ectopic innervation of outer hair cells by a subset of type I SGNs (Defourny, Poirrier et al. 2013). EPHA4 interacts with EFNA5 to control target specification, expressed primarily in OHCs and a subset of type I SGNs. Previously, *Efna5* was reported to be expressed in a subset of type I SGNs, but recent evidence has implicated Efna5 primarily in type II SGNs (Petitpre, Wu et al. 2018, Sanders and Kelley 2022). The otic mesenchyme surrounding the SGN axons also expresses *EphA4* (Coate, Spita et al. 2015). This pattern of EPHA4 expression guides fasciculation through EFNB2 interactions in the developing SGN axons. Another *Eph* receptor, *EphA7*, in SGNs, is required for neurite outgrowth. Loss of *EphA7* resulted in more sparse fiber bundles and fewer synaptic contacts on inner hair cells (Kim, Ibrahim et al. 2016). *EfnB2* is one of the ephrin ligands, and a high level of *Efnb2* mRNA expression was detected during SGN development, especially at E13.5 and E15.5, when SGN fasciculation initiates. Deletion of *Efnb2* in SGNs showed defective fasciculation of inner radial bundles and an increased number of aberrantly crossing SGN axons between fascicles (Micucci, Sperry et al. 2015). These studies implicate ephrin signaling in the development of peripheral auditory circuits.

The axon guidance deficits observed in the *Chd4* cKO cochleae are distinct from the Eph/ephrin studies. Using *Ngn1* CreER^T2^, we perturbed only SGNs, leaving other cell types, such as hair cells and otic mesenchyme, untouched. Instead of inactivating the function of ephrin receptors or ephrin ligands, *Chd4* deletion increased transcript levels of *Epha4*, *Epha7*, and *Efnb2* in SGNs, revealing a molecular perturbation relevant to CHD4-dependent neurodevelopmental processes. The *Chd4* cKO model also differs from single knockouts of Eph receptors and Ephrin ligands in that it affects the expression of multiple axon guidance genes. The subtle cellular phenotypes in single *Eph* and *Efn* knockout mice suggest that the proper wiring of the peripheral auditory circuit may require the combined effects of multiple *Eph* and *Efn* members. Finally, the *Chd4* cKO model differs from ablation of the Eph and Efn genes in that it increases transcription of its target genes. Our findings align with the involvement of numerous *Eph* and *Efn* genes during SGN development and suggest that CHD4 may coordinate the expression levels of several *Eph* and *Efn* to shape the type I and II SGN innervation patterns.

Congenital hearing loss syndromes are often associated with pathogenic genetic variants in a single gene. Hearing loss can be syndromic or non-syndromic. Syndromic hearing impairment is associated with clinical features in other organ systems, while non-syndromic hearing impairment has no discernible clinical abnormalities other than in the middle or inner ear. Syndromic hearing loss encompasses over 400 genetic conditions (Toriello, Reardon et al. 2004). The characteristics of syndromic hearing loss vary across syndromes and can affect hearing in one or both ears. The extent of hearing loss varies widely, ranging from profound to mild. Different syndromes exhibit distinct patterns of hearing loss. Improper development or dysfunction of the sensory cells and neurons of the inner ear correlates with many forms of syndromic hearing loss. Patients with Sifrim-Hitz-Weiss disease show variable congenital disabilities. This observation is reminiscent of CHARGE syndrome, a genetic disorder caused by pathogenic variants in CHD7. CHD7 and CHD4 are part of the chromodomain helicase DNA-binding protein family (Micucci, Sperry et al. 2015). Global chromatin regulators such as CHD7 and CHD4 can robustly affect transcription and trigger cell-autonomous changes during critical periods in development. We show that altering the epigenetic landscape by deleting CHD4 early in otic development (E8.5-E10.5) affects the pathfinding and connectivity of SGN peripheral axons at multiple steps of axon guidance, contributing to hearing loss in SIHIWES.

We believe that cellular heterogeneity and increasing variance are central aspects caused by mutations in chromodomain helicase DNA-binding proteins. There are four distinct subtypes of SGNs: type Ia, Ib, Ic, and type II (Petitpre, Wu et al. 2018, Shrestha, Chia et al. 2018, Sun, Wang et al. 2022). CHD4 may function differently across neuronal subtypes, resulting in distinct cellular phenotypes. The difference is evident in the axon-guidance phenotypes observed in type I versus type II neurons. Ablation of CHD4 affected the fasciculation of radial fibers and branching in the inner spiral plexus region for type I SGNs. In contrast, type II neurons displayed a circuitous path to innervate the base of outer hair cells and inappropriate turning at the outer hair cell region. Deleting CHD4 affected the peripheral axons differently in type I and II SGNs.

In addition to heterogeneity among neuronal subtypes, variability was observed within type I and II fibers. The phenotypic variability includes the extent of type I radial fiber bundling and branching of terminals in the inner spiral plexus. CHD4 function in SGN type Ia, b, and c fibers may contribute to the observed variability. Cellular heterogeneity, however, cannot be attributed to the improper turning of some type II peripheral axons. Some type II fibers are seemingly unaffected, whereas others turn in the opposite direction. Stochasticity in molecular processes may be a contributing mechanism. Stochastic gene expression involves biochemical processes such as transcription and translation. The limited number of transcription factors leads to biological variability, as evidenced by dramatic differences in transcript levels even within isogenic cells (Raj and van Oudenaarden 2008). Deletion of CHD4 may alter the chromatin state of cis-regulatory elements and impact the stochasticity in transcription. The epigenetic perturbation could increase both transcript levels and the variance of specific genes following CHD4 ablation. We observed an increase in the median numbers of *Epha4*, *Epha7*, and *Efnb2* transcripts in *Calb2*-marked SGNs, concomitant with an increase in the variance of transcript numbers. The increase in transcripts and variance was specific for *Epha4, Epha7, and Efnb2*, not *Calb2*. The larger variance results in a broader distribution of neurons with *Epha4, Epha7, and Efnb2* and displays a cell population that exceeds the normal range of *Epha4, Epha7, and Efnb2* levels in control SGNs. The percentage of neurons with increased *Epha4, Epha7, and Efnb2* transcripts may possess altered protein expression levels of EPHA4, EPHA7, and EFNB2 and perturb axon guidance. In contrast, cells that retain *Epha4, Epha7, and Efnb2* transcripts within the normal range may retain normal axon pathfinding. The increase in transcriptional variance may contribute to the heterogeneous cellular phenotypes observed for type I and type II SGNs.

In summary, multiple axon guidance deficits of type I and II SGNs were observed using a *Chd4* cKO mouse model. These deficits likely arise at different developmental time points and implicate CHD4 function at varying steps of SGN pathfinding. The candidate CHD4 binding sites were identified at promoters and enhancers that regulate the transcription of a subset of *Eph* receptors and *Efn* ligands. CHD4, as part of the core NuRD complex, may impose a repressive epigenetic state on the cis-regulatory elements of these genes, attenuating transcription until the appropriate time during development. Inactivation of CHD4 increases transcript levels of a subset of axon guidance genes, including Eph receptors and ephrin ligands. Beyond this, our CHD4 occupancy motif analysis suggests that CHD4 directly targets additional guidance molecules, including members of the semaphorin, plexin, and frizzled families. While the precise transcript dynamics of these families remain to be characterized in future expression studies, their direct genomic targeting implies a broader epigenetic network through which CHD4 orchestrates the stereotypic peripheral axon patterning in the cochlea. Together, these results support a model in which CHD4-mediated chromatin remodeling fine-tunes transcription of multiple axon guidance genes, enabling precise regulation and also the formation of neuronal circuits during development.

## Accession Numbers

The GEO accession number for the CUT&Tag raw fastq data, along with the processed bigwig and SEACR peak files, is GSE250033. IGV session with the CUT&Tag tracks can be provided upon request.

## Author contribution

JK and EM designed the study and executed the experiments, including animal procedures, tissue collection and processing, cell cultures, immunofluorescence and RNAscope studies, image acquisition, and data analysis. JN performed CUT&Tag, and JQ analyzed the CUT&Tag data. AL and AMH performed ABR, and KYK analyzed the ABR data. KYK conceived the project, performed motif analysis, and assisted with CUT&Tag and RNAscope analyses. JK, EM, and KYK wrote the manuscript.

## Acknowledgments

We thank Dr. Katia Georgopoulos (Mass General Hospital) and Dr. Lisa Goodrich (Harvard Medical School) for providing the *Chd4*^flox/flox^ animals and advice on dosing the *Ngn1* CreER^T2^ animals, respectively. We thank Dr. Zainab Tanvir at the SAS-HGINJ imaging facility for her technical expertise. NIH R01 DC015000 and DC022483, which supported this work for KYK.

## Supplemental Figure Legend

**Supplemental Figure 1.**
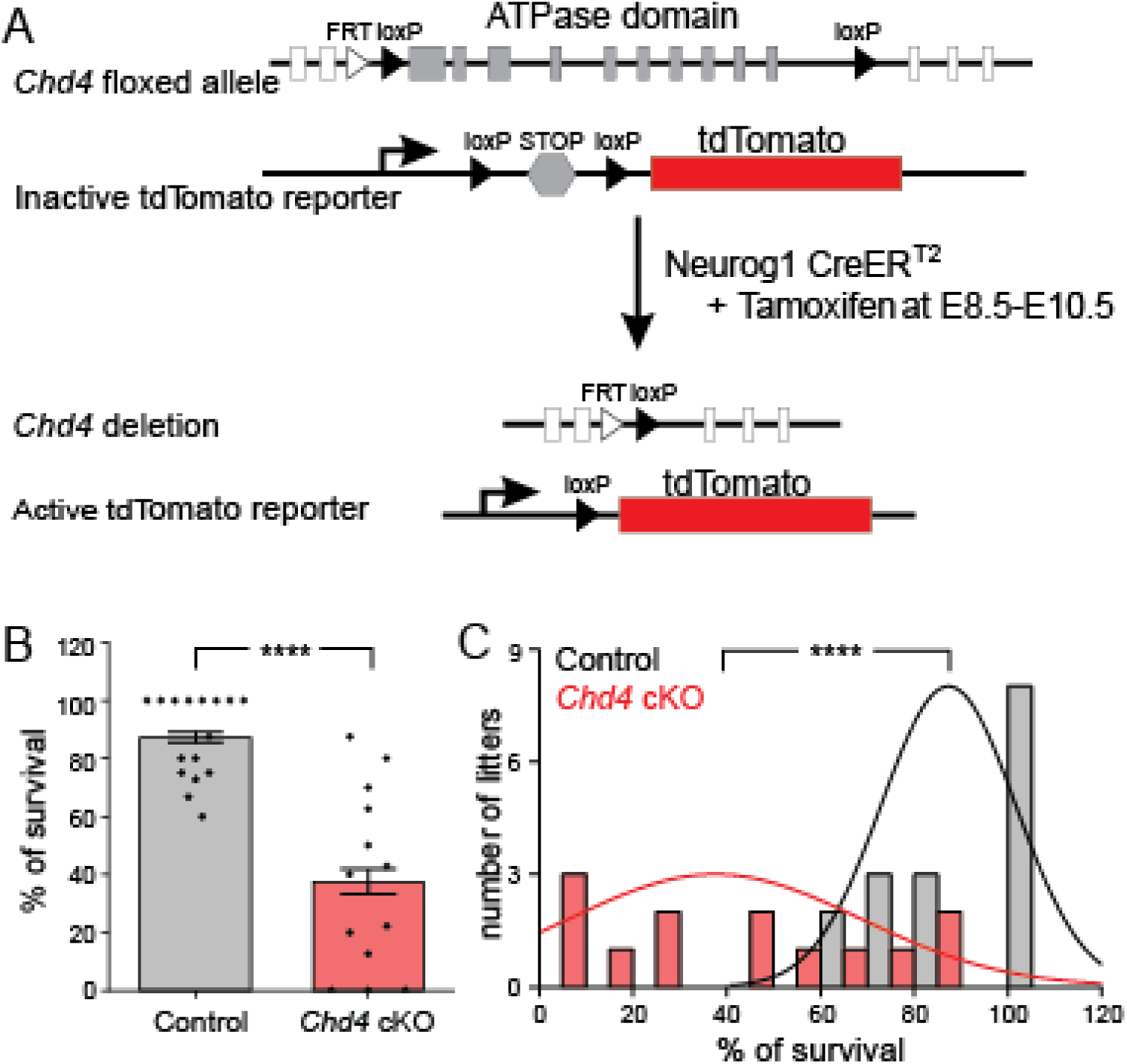
Survival of *Chd4* cKO mice. (A) Schematic illustration of Cre-mediated *Chd4* deletion following tamoxifen induction in *Chd4* cKO mice containing *Ngn1* CreER^T2^; *Chd4*^flox/flox^; tdTomato alleles. Administration of tamoxifen at E8.5-E10.5 deletes loxP-flanked exons encoding the *Chd4* ATPase domain and activates tdTomato reporter expression. Animals without the *Chd4* cKO allele (*Ngn1* CreER^T2^; tdTomato) were used as controls. (B) Average percentage of viable pups from all collected control and mutant litters (p < 4.36 × 10^-6^; control, n=16, and *Chd4* cKO, n=13 litters). (C) Distribution of the percentages of surviving E18.5 pups from control and *Chd4* cKO litters after tamoxifen treatment. The p-values and sample size are listed. The Student’s *t*-test was used for statistical analysis.

**Supplemental Figure 2.**
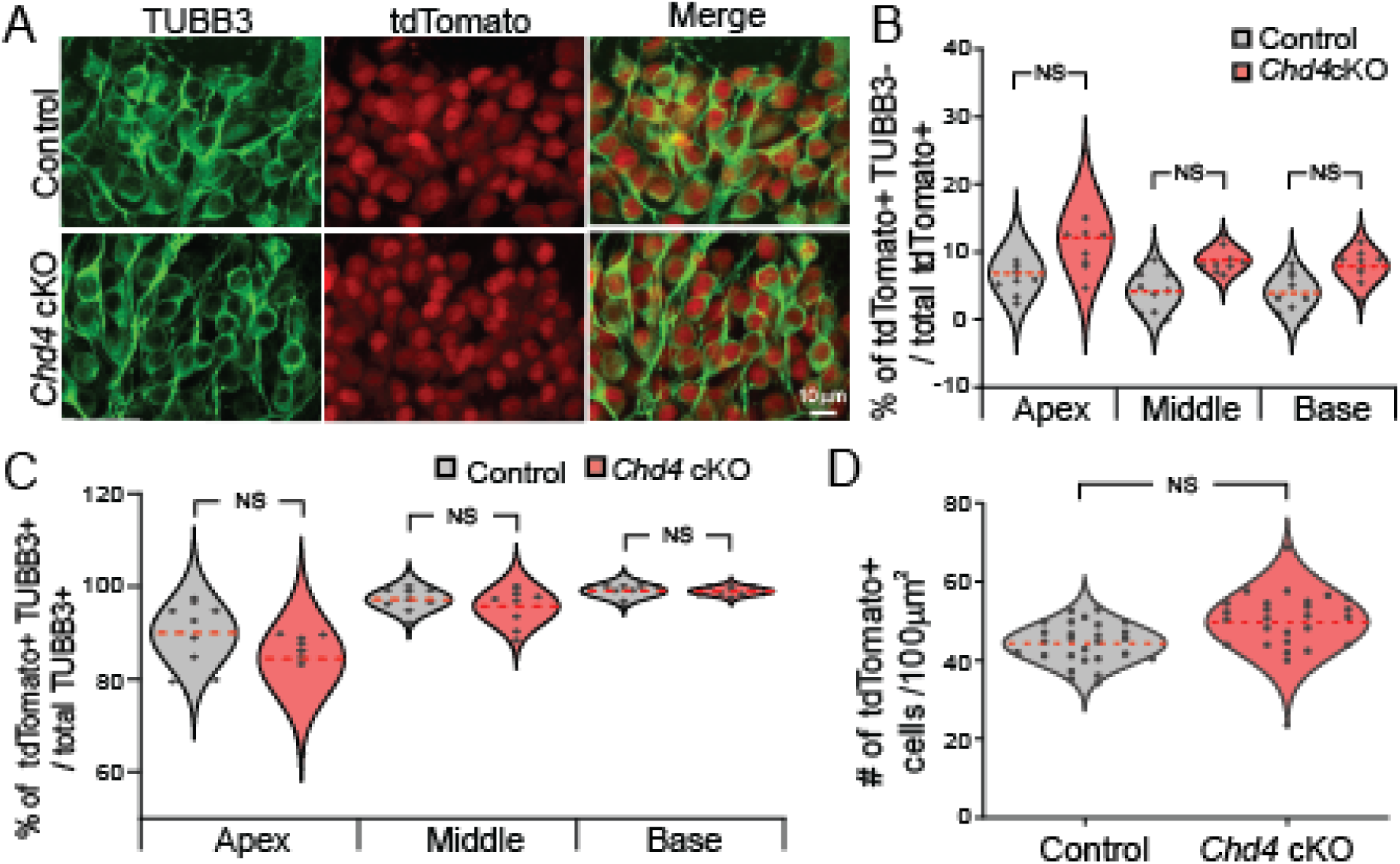
CHD4 is not required for SGN viability. (A) Confocal micrographs from the spiral ganglion. Yellow arrows indicate tdTomato^+^ and TUBB3^-^ cells. (B) Percentages of tdTomato^+^ and TUBB3^-^ cells found within the apex, middle, and base regions of the cochlea (apex, p < 0.136, middle, p < 0.095; base, p < 0.148; control, n=3 and *Chd4* cKO, n=3 cochlea) (C) Percentages of cells expressing tdTomato and TUBB3 from the base to apex of the cochlea from control and *Chd4* cKO cochleae (apex p < 0.30; middle p < 0.60; base, p < 0.90; control, n=3 and *Chd4* cKO, n=3 cochleae) (D) Normalized number of tdTomato^+^ cells per 100 µm^2^ area in control and *Chd4* cKO cochleae (p < 0.08; control, n=27 and *Chd4* cKO n=27 images were analyzed from control, n=3 and *Chd4* cKO, n=3 cochleae). Student’s *t*-test was used for statistical analysis based on the number of cochleae. Scale bar as marked.

**Supplemental Figure 3.**
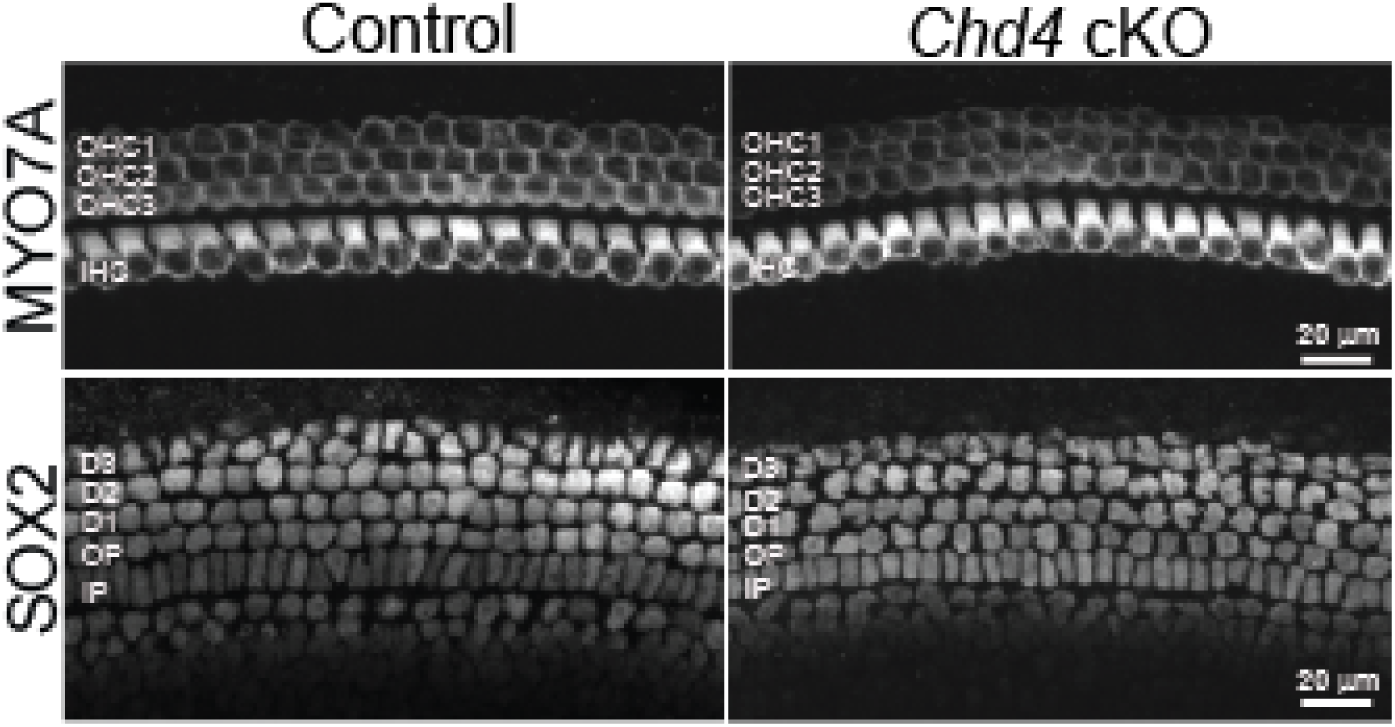
The arrangement of supporting and hair cells is unaffected following the deletion of *Chd4* within spiral ganglion neurons. Immunofluorescent labeling of MYO7A and SOX2 marked hair cells and supporting cells from E18.5 cochleae. Confocal stacks from control and *Chd4* cKO cochleae with MYO7A-marked hair cells and SOX2-marked supporting cells. OHC (outer hair cell), IHC (inner hair cell), D (Deiters’ cells), OP (outer phalangeal cells), and IP (inner phalangeal cells). Scale bar as marked.

**Supplemental Figure 4.**
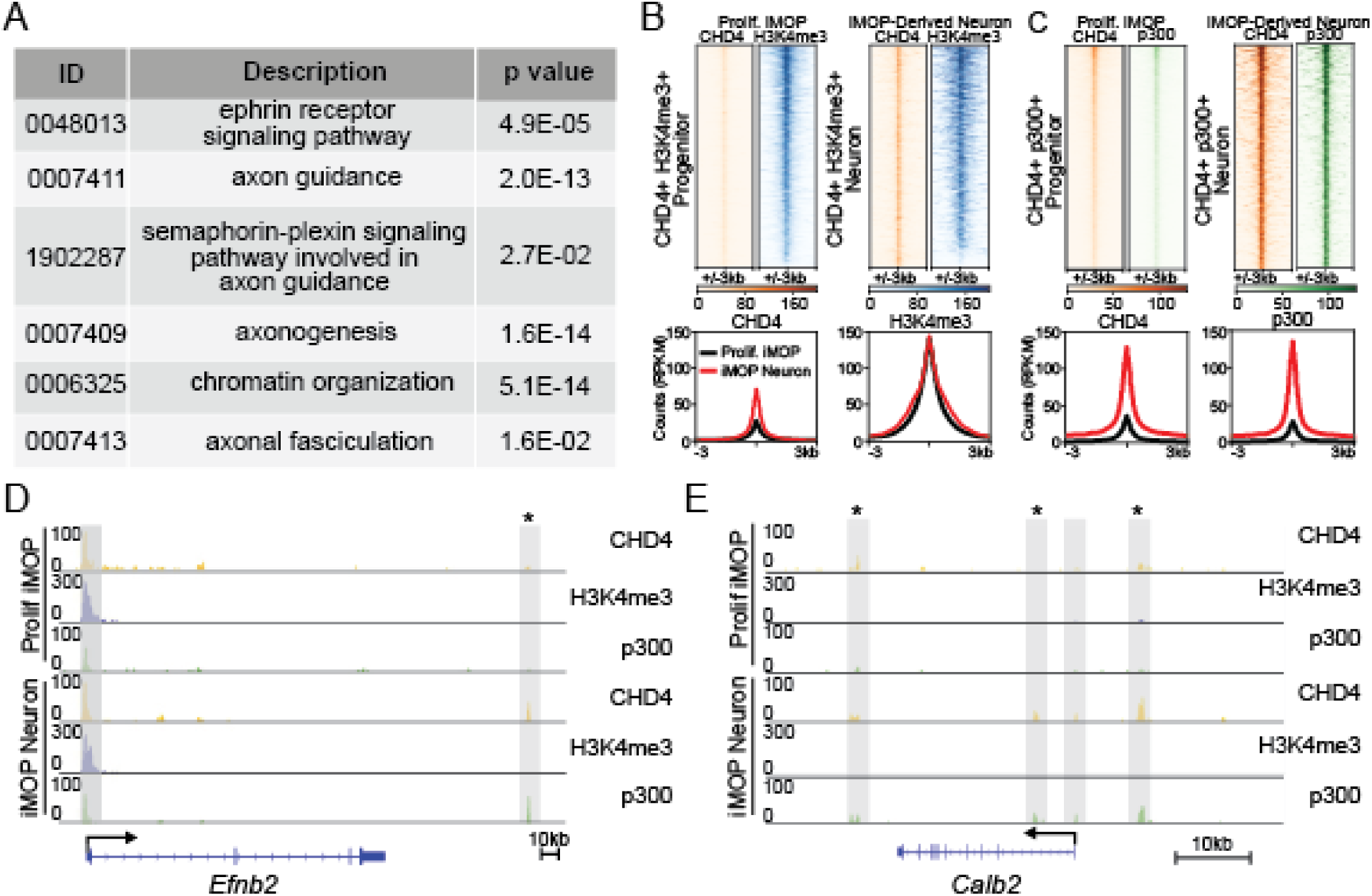
Identifying CHD4 binding at promoters and enhancers of target genes. (A) Table showing Gene Ontology ID, description of each category, and corresponding adjusted p-values using the Benjamini-Hochberg method for multiple testing correction. (B) Heatmaps and profile plots of CHD4 and H3K4me3 signals within ± 3kb of the summit regions of CHD4+ H3K4me3+ in both proliferating iMOP and iMOP-derived neurons. (C) Heatmaps and profile plots of CHD4 and p300 signals within ± 3kb of the summit regions of CHD4+ p300+ in proliferating iMOPs and iMOP-derived neurons. (D) Enrichment of CHD4 (orange), H3K4me3 (blue), and p300 (green) at the specified genomic regions of *Efnb2* in proliferating iMOPs and iMOP-derived neurons. Highlighted genomic regions represented H3K4me3-marked promoters, while highlighted regions with asterisks denote regions with increased CHD4 and p300 in iMOP-derived neurons. The number of asterisks indicates the number of peaks that show increased CHD4 and p300 levels. Scale bars as marked.

**Supplemental Figure 5.**
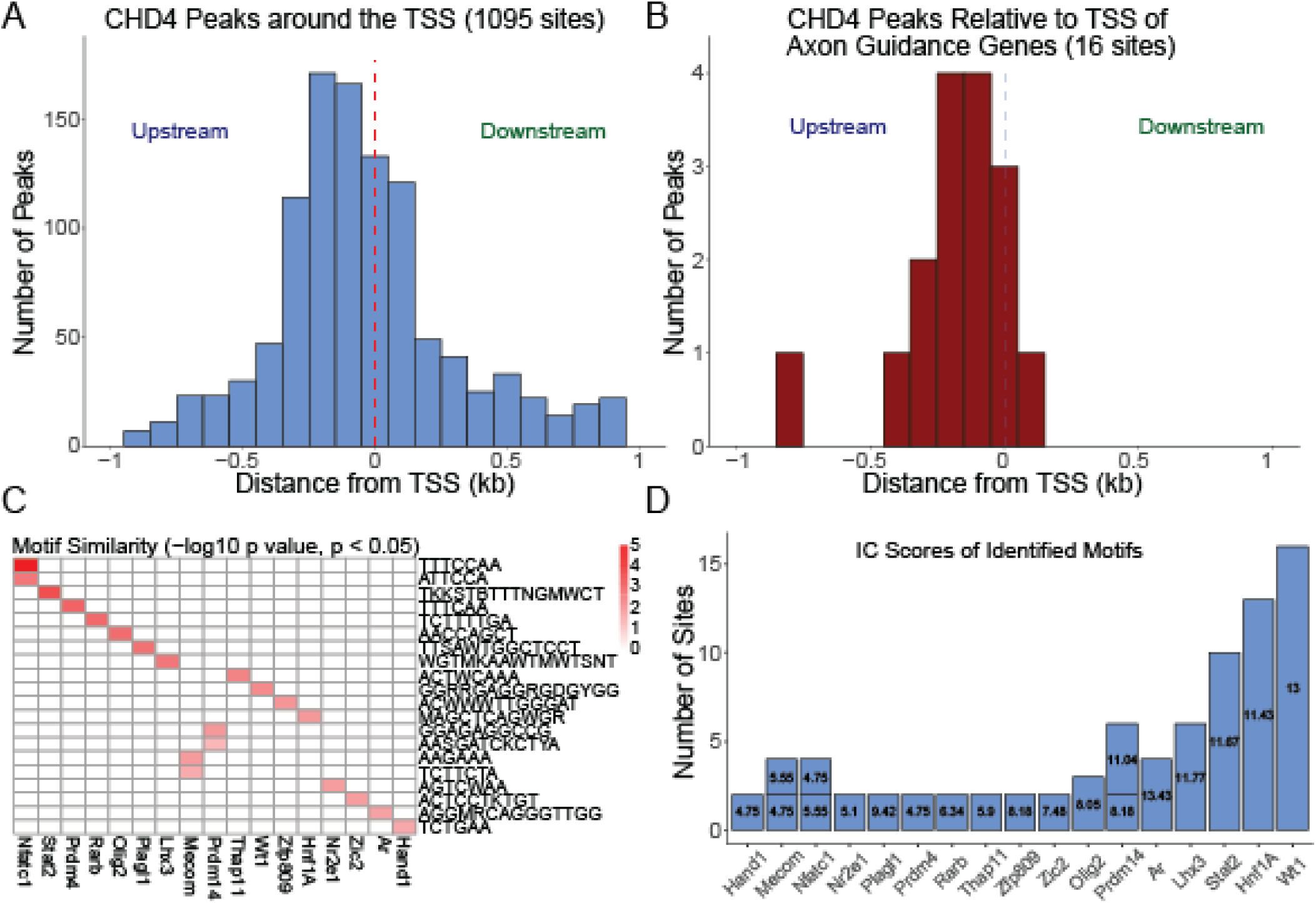
*De novo* motif discovery and annotation. (A) A histogram displaying 1,095 CHD4 peaks relative to the transcriptional start sites (TSS) from all the CHD4 binding sites. The dashed red line separates the upstream and downstream regions at the TSS. (B) Distribution of 16 CHD4 peaks relative to TSS at axon guidance genes. (C) A heatmap depicting the pairwise comparison between a set of *de novo* motif sequences with known transcription factor (TF) names. The consensus sequences are shown on the right, and sites corresponding to TF binding sites are at the bottom. The color intensity represents the statistical significance similarity comparisons (p < 0.05). (D) Information Content (IC) scores of annotated motifs showing the number of occurrences with the associated Information Content (IC) score. The IC score reflects the specificity or conservation of the motif.

**Supplemental Figure 6.**
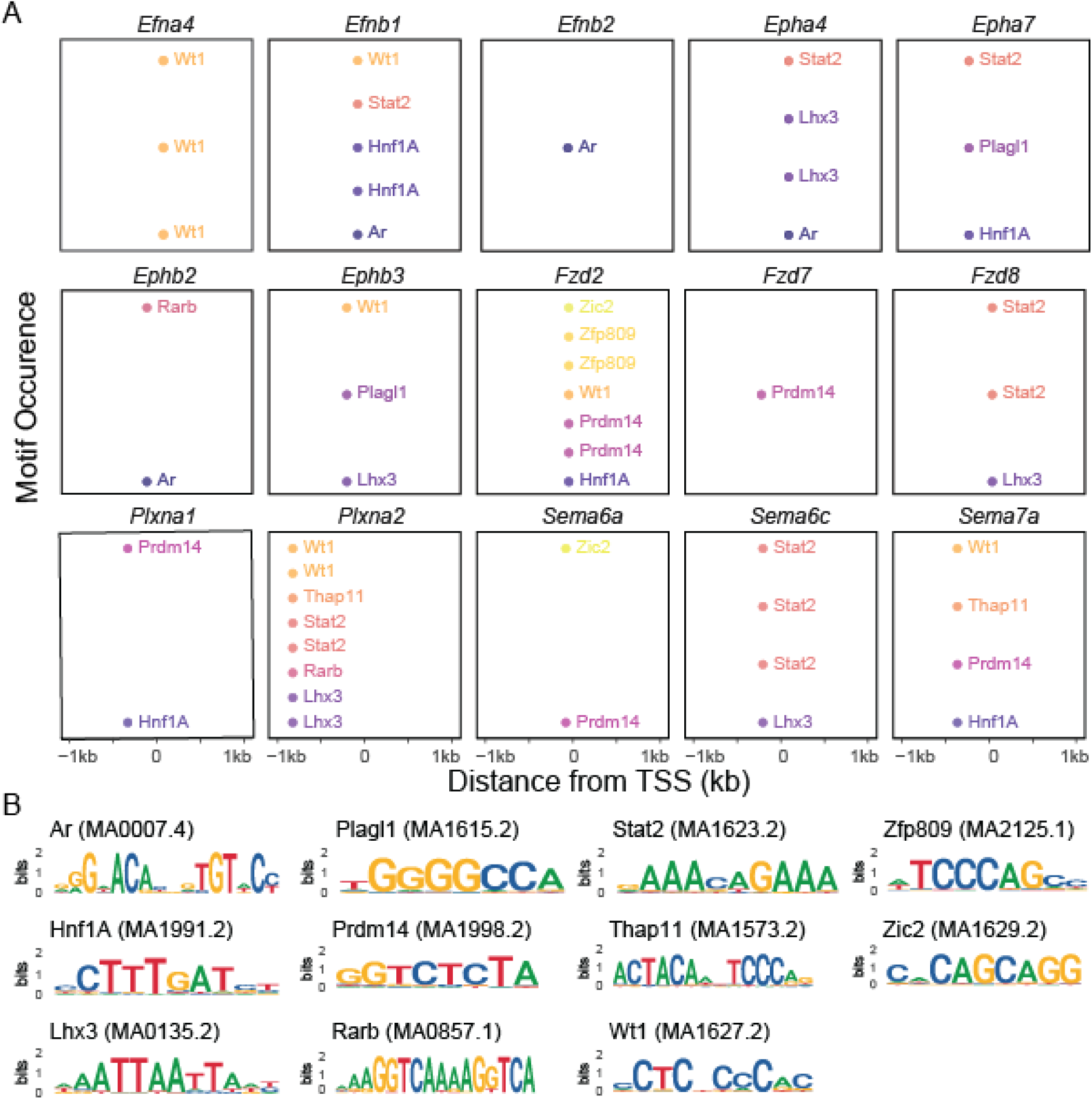
Motif consensus sequences and occurrence. (A) The occurrence and position relative to the TSS of annotated motifs. Each panel represents the genomic region around the TSS of the axon guidance gene. Within each panel, dots represent the occurrence of a TF motif relative to the TSS. (B) Sequence logos for the various TF binding motifs identified in the axon guidance genes. Each logo contains the motif name and its unique JASPAR accession number.

